# Muscle-Specific Pyruvate Kinase Isoforms, Pkm1 and Pkm2, Regulate Mammalian SWI/SNF Proteins and Histone 3 Phosphorylation During Myoblast Differentiation

**DOI:** 10.1101/2024.04.10.588959

**Authors:** Monserrat Olea-Flores, Tapan Sharma, Odette Verdejo-Torres, Imaru DiBartolomeo, Paul R. Thompson, Teresita Padilla-Benavides, Anthony N. Imbalzano

## Abstract

Pyruvate kinase is a glycolytic enzyme that converts phosphoenolpyruvate and ADP into pyruvate and ATP. There are two genes that encode pyruvate kinase in vertebrates; *Pkm* and *Pkl* encode muscle- and liver/erythrocyte-specific forms, respectively. Each gene encodes two isoenzymes due to alternative splicing. Both muscle-specific enzymes, Pkm1 and Pkm2, function in glycolysis, but Pkm2 also has been implicated in gene regulation due to its ability to phosphorylate histone 3 threonine 11 (H3T11) in cancer cells. Here, we examined the roles of Pkm1 and Pkm2 during myoblast differentiation. RNA-seq analysis revealed that Pkm2 promotes the expression of *Dpf2/Baf45d* and *Baf250a/Arid1A*. Dpf2 and Baf250a are subunits that identify a specific sub-family of the mammalian SWI/SNF (mSWI/SNF) of chromatin remodeling enzymes that is required for activation of myogenic gene expression during differentiation. Pkm2 also mediated the incorporation of Dpf2 and Baf250a into the regulatory sequences controlling myogenic gene expression. Pkm1 did not affect expression but was required for nuclear localization of Dpf2. Additionally, Pkm2 was required not only for the incorporation of phosphorylated H3T11 in myogenic promoters, but also for the incorporation of phosphorylated H3T6 and H3T45 at myogenic promoters via regulation of AKT and protein kinase C isoforms that phosphorylate those amino acids. Our results identify multiple unique roles for Pkm2 and a novel function for Pkm1 in gene expression and chromatin regulation during myoblast differentiation.

## INTRODUCTION

The formation of skeletal muscle is a complex process that originates in the embryonic mesoderm and results in the formation of functional muscle tissue by birth. Embedded in the skeletal muscle are stem cells called satellite cells that will be responsible for post-natal myogenesis. The development of skeletal muscle tissue in both embryos and post-natal organisms involves the development of myoblasts from myogenic precursors and subsequent differentiation into multi-nuclear myotubes that ultimately form contractile myofibers (1–5). The myoblast to myotube transition has been well-studied as it is amenable to replication in cell culture (6, 7) and has provided numerous conceptual advances in the understanding of gene regulation by sequence-specific DNA-binding transcription factors (6, 8, 9) and co-regulatory factors, many of which have been identified as histone modifiers and chromatin remodeling enzymes (10–13).

Integration of the mechanisms by which this multitude of factors cooperates to regulate myoblast proliferation and the onset and the execution of the myoblast differentiation program is complicated by the impact of developmentally relevant signaling molecules and their effector molecules. These effector molecules, including protein kinases and phosphatases, usually have multiple substrates, making a comprehensive understanding of the myoblast to myotube transition difficult (14–16). Nevertheless, mechanistic investigation of known and suspected players in the process usually provides novel insights as well as additional questions about overall regulation of differentiation.

Our lab’s efforts have for years concentrated on the functional properties of the ATP-dependent chromatin remodeling enzymes of the mSWI/SNF family. The mSWI/SNF enzymes utilize the energy obtained from ATP hydrolysis to transiently alter the chromatin configuration on target DNA sequences, which can result in nucleosome sliding or eviction and the stable binding or exclusion of transcription or other regulatory factors (17–22). These enzymes alter local myogenic gene chromatin structure to facilitate the assembly of active transcription complexes that initiate or up-regulate the expression of differentiation-specific genes (11, 23, 24). In addition to working within the context of activities by multiple histone modifying enzymes, the mSWI/SNF enzymes themselves are regulated by signaling pathways that involve modification by p38, PKC and calcineurin, and CK2, amongst others (15, 24–28).

Here, we investigated the roles of the muscle-specific pyruvate kinase enzymes in the regulation of chromatin remodeling enzymes and chromatin modifications during myoblast proliferation and differentiation. Pyruvate kinase (PK) catalyzes the final rate-limiting step of glycolysis by converting phosphoenolpyruvate and ADP to pyruvate and ATP (29). Alternative splicing of two pyruvate kinase genes, the liver/erythrocyte-specific enzyme (*Pklr*) and muscle-specific enzyme *Pkm*, generates four isoenzymes in mammalian cells (30, 31). The mutually exclusive splicing of exon 9 or 10 of the *Pkm* pre-mRNA produces *Pkm1* or *Pkm2* (30, 32). An isoform switch from *Pkm2* to *Pkm1* occurs between late embryonic stages and early post-natal development of skeletal muscle (33–37). Given the requirement for the mSWI/SNF chromatin remodeling enzyme complexes in myoblast differentiation and the increasing evidence for multiple enzymes controlling the phosphorylation state of mSWI/SNF subunits, we asked whether there might be a functional connection between Pkm and mSWI/SNF enzymes.

While there is little known about Pkm and chromatin remodeling enzymes, there is a definite connection between Pkm enzymes and the phosphorylation of H3T11. PKM2 had been identified as a regulator of cell proliferation through undetermined mechanisms (38–41). Subsequent work suggested a correlation between PKM2 enzymatic activity and the dissociation of histone deacetylases and acetylation of H3 at cell cycle-related gene promoters upon EGFR-stimulation (42). Pkm2, but not Pkm1, was then shown to phosphorylate H3T11 but not H3T3 or T6 (43). H3T11 phosphorylation served as the triggering event that precipitated HDAC dissociation and subsequent molecular events associated with gene activation (43). The data document the idea of a non-metabolic role for Pkm2 in regulating gene expression. Further support for such a role can be found in the yeast literature. PYK1, a yeast homolog of PKM2, phosphorylates H3T11 as a part of the protein serine-responsive SAM-containing metabolic enzyme (SESAME) complex that links H3K4 methylation and H3T11 phosphorylation in response to glycolysis and serine metabolism (44, 45). SESAME complex dependent phosphorylation of H3T11 also represses transcription of genes near telomeres by preventing the nuclear export and degradation of the SIR2 repressor (45).

Like H3T11 phosphorylation, H3T6 phosphorylation can be associated with active gene expression. Protein kinase C beta (PKCβ) phosphorylates H3T6 to prevent lysine specific demethylase 1 (LSD1) from demethylating H3K4, thereby promoting and/or maintaining gene expression (46). The PKC delta isoform (PKCδ) was also identified as the relevant kinase for H3T45 phosphorylation in vivo and in vitro during apoptosis (47). Consistent with this result, PKC1 from *S. cerevisiae* phosphorylates H3T45, which is required for subsequent H3K56 acetylation (48). Other work, however, implicated AKT (protein kinase B) as an H3T45 kinase in response to DNA damage (49). Though the highest concentration of phosphorylated H3T45 was located around the site of transcriptional termination at affected genes, signal was located throughout the targeted loci. Given this information and our prior efforts in characterizing PKCβ and calcineurin working in opposition to regulate the phosphorylation of the Brg1 ATPase of the mSWI/SNF enzymes during myoblast differentiation (25, 28), we included phosphorylation of H3T6 and H3T45 in our study.

Here we report that both Pkm1 and Pkm2 have functions related to gene expression that appear to be independent of the roles of these enzymes in glycolysis. Pkm2, but not Pkm1 regulated the expression of two mSWI/SNF chromatin remodeling enzyme subunits, Dpf2 and Baf250a, that belong to the sub-family of mSWI/SNF enzymes previously identified as mediating the expression of myogenic genes during myoblast differentiation (50, 51). Pkm1, but not Pkm2, contributed to nuclear localization of the Dpf2 subunit. In addition, Pkm2 mediated the incorporation of not just phosphorylated H3T11, a known substrate of Pkm2, into myogenic gene regulatory sequences but also incorporation of phosphorylated H3T6 and phosphorylated H3T45, likely via indirect mechanisms.

## MATERIALS AND METHODS

### Antibodies

Hybridoma supernatant against myosin heavy chain (MHC; MF20, deposited by D. A. Fischman) was obtained from the Developmental Studies Hybridoma Bank (University of Iowa, by A. Kawakami). Mouse anti-Brg1 (G-7; sc-17796), anti-Lamin A/C (4A58) and alpha Tubulin (B-7) were obtained from Santa Cruz Biotechnologies (Dallas, TX, USA). Rabbit anti-phospho-Histone H3T6 (AP0093), -Phospho-Histone H3T11 (AP0093), and -Phospho-Histone H3T45 (AP0898) were from Abclonal Technologies (Woburn, MA, USA). Rabbit anti-Pkm1 (D30G6), -Pkm2 (D78A4), Arid1a/Baf250a (D2A8U), -vinculin (E1E9V), and Dpf2 (E7N8J) were from Cell Signaling Technologies (Danvers, MA, USA). Secondary antibodies used for western blot were HRP-conjugated anti-mouse and anti-rabbit (31430 and 31460, respectively) and for immunofluorescence were the goat anti-rabbit IgG Alexa Fluor Plus 488 (A11008) from Thermo Fisher Scientific (Waltham, MA, USA).

### Cell culture

The C2C12 established cell line is a subclone of the murine myoblast line established from normal adult C3H mouse leg muscle and has been used extensively as a model for skeletal muscle differentiation *in vitro* (7, 52). The HEK293T cell line is a derivative human embryonic kidney cell line that expresses a mutant version of the SV40 large T antigen and was used for lentiviral production. C2C12 and HEK293T cells were purchased from the American Type Culture Collection (ATCC, Manassas, VA) and were maintained at sub-confluent densities in proliferation media containing Dulbecco’s modified Eagle’s medium (DMEM) supplemented with 10% Fetal Bovine Serum (FBS, Gibco), 1% penicillin-streptomycin in a humidified incubator at 37 °C with 5% CO_2_. To induce myoblast differentiation, C2C12 cells cultured to 80% confluence were incubated in DMEM media supplemented with 2% Horse Serum (HS, Gibco), Insulin-Transferrin-Selenium-A (Gibco) and 1% penicillin-streptomycin in a humidified incubator at 37 °C with 5% CO_2_. The specific chemical inhibitor of PKM2 activity, 3K (Selleck USA, Houston, TX, United States) (53), was used at 2 µM following the initial titration experiments.

### Lentivirus production for shRNA transduction of C2C12 cells

Mission plasmids encoding a control scrambled shRNA (Scr) and individual shRNA oligos against *Pkm1* and *Pkm2* (**Supp. Table 1**) were cloned in the pLKO_TRC001 vector (Addgene cat no. 10878) as previously described (54). Plasmids were transformed into TOP10 chemically competent cells, and isolated with the pure yield plasmid miniprep system (Zymo Research) following the manufacturer’s instructions. The purified plasmids encoding for the specific shRNAs were verified by Sanger sequencing (Quintara Biosciences, Cambridge, MA, USA).

### Gene expression analysis

RNA was isolated from three independent biological replicates of differentiating C2C12 myoblasts expressing sh*Scr*, sh*Pkm1* (shRNA#2), or sh*Pkm2* (shRNA#2) with TRIzol (Invitrogen) following the manufacturer’s instructions. cDNA synthesis was performed with 1 μg of RNA as template for reverse transcription and the HiFiScript gDNA Removal RT MasterMix Kit (Cwbio), following the manufacturer’s protocol. Quantitative PCR was performed with Fast SYBR green master mix on the ABI StepOne Plus Sequence Detection System using the primers listed in **Supp. Table 2**, and the delta threshold cycle value (2^ΔΔCT^; (55)) was calculated for each gene and represented the difference between the CT value of the gene of interest and that of the control gene, *Eef1a1α*.

### Subcellular fractionation

Cell fractionation was performed following the rapid, efficient, and practical (REAP) method (56). Briefly, differentiated myoblasts were washed three times with ice-cold 1X PBS and resuspended in 1 ml of PBS. The samples were centrifuged, the supernatant was discarded, and the cell pellet was resuspended in 900 μl of ice-cold PBS containing 0.1% NP-40. Cells were resuspended for 40 times with a P1000 micropipette tip to release the nuclei. A portion of this cell suspension (300 μL) was separated (whole cell lysate) and the remaining fractions were centrifuged at 10,000 rpm. The supernatant, or cytosolic fraction, was recovered in a new tube, and the nuclei-containing pellet was washed once more with 1 mL of PBS containing 0.1% NP-40. The nuclear fraction was again centrifuged at 10,000 rpm, the supernatant was discarded, and the pellet was resuspended in 150 μL of PBS with 0.1% NP-40.

### Western blot analyses

Differentiating C2C12 (control and knockdown) myoblasts were washed three times with ice-cold PBS and solubilized with RIPA buffer (10 mM piperazine-N,N-bis(2-ethanesulfonic acid) pH 7.4, 150 mM NaCl, 2 mM ethylenediamine-tetraacetic acid (EDTA), 1% Triton X-100, 0.5% sodium deoxycholate, and 10% glycerol) containing protease inhibitor cocktail (Thermo Fisher Scientific). Protein content was determined by Bradford analysis (57), and 25 μg of each sample were prepared for SDS-PAGE by boiling in Laemmli buffer. The resolved proteins were electro-transferred to PVDF membranes (Millipore). The proteins of interest were detected with the specific antibodies indicated in the figure legends, followed by species-specific peroxidase-conjugated secondary antibodies and Tanon chemiluminescence detection system (Abclonal Technologies). Densitometric analyses were performed with ImageJ software v. 1.8.

### Immunocytochemistry

Differentiating C2C12 myoblasts (control and knockdown) were fixed overnight with 10% formalin-PBS at 4 °C. Cells were washed three times with PBS and permeabilized for 30 min with PBS containing 0.2% Triton X-100. Immunocytochemistry was performed on differentiating myoblasts using the myogenin or MHC hybridoma supernatant and was developed with the universal ABC kit (Vector Labs) following the manufacturer’s protocol. Images were acquired with an Echo Rebel microscope using the 10X objective (Echo).

### Determination of fusion index

The fusion index was calculated as the percentage of nuclei/cells stained with myosin heavy chain as compared to the total number of nuclei/cells. Edges and regions that did not show good adhesion were discarded from the analysis. Three independent biological replicates were grown in 24-well plates, and cells were induced to differentiate as described above. Quantification was performed with ImageJ software v. 1.8.

### Immunofluorescence

Control and knockdown differentiating (48 h) C2C12 myoblasts were grown on cover slides and fixed overnight in 10% formalin-PBS at 4 °C. The monolayers were washed with PBS, permeabilized with 0.2% Triton X-100 in PBS for 15 min, incubated for 1 h at RT in blocking solution containing PBS, 0.2% Triton X-100, 3% FBS. Samples were incubated overnight with anti-Dpf2 antibodies in blocking buffer at 4 °C. The next day, the cells were incubated for 3 h at RT with a goat anti-rabbit Alexa-488 secondary antibody in blocking solution and for 30 min with DAPI (F6057, SigmaAldrich, St. Louis, MO, United States). Microscopy and image processing were performed using a Leica SP8 Confocal Microscope and the Leica Application Suite X (Leica Microsystems Inc., Buffalo Grove, IL, United States).

### Chromatin immunoprecipitation assays

Chromatin immunoprecipitation (ChIP) assays for C2C12 cells were performed in triplicate from three independent biological samples as previously described (51, 58, 59). Briefly, chromatin digestion was performed using a kit (ChIP kit #9005, Cell Signaling Technologies, Danvers, MA, USA), following the manufacturer’s instructions, then crosslinked lysates of myoblasts were incubated overnight with anti-phospho-H3T6, phospho-H3T11, phospho-H3T45, Baf250a, Dpf2, total H3 or normal rabbit IgG. Crosslinking was reversed, and DNA was purified using ChIP DNA Clean and Concentrator Columns (Zymo Research) following the manufacturer’s instructions. qPCR was performed using Fast SYBR green master mix on the ABI StepOne Plus Sequence Detection System. Primers used are listed in **Supp. Table 2**. Quantification was done using the comparative Ct method (ΔCT; (55)) to obtain the percent of total input DNA pulled down by each antibody.

### Statistical analysis

Statistical analyses were performed using Graph Pad Prism 7.0b. Multiple data point comparisons and statistical significance were determined using one-way analysis of variance (ANOVA) and the comparisons were performed using Dunnett’s multiple comparison tests. Experiments where *p< 0.05* were considered statistically significant.

## RESULTS

### *Pkm2* knockdown or Pkm2 inhibition affected myoblast proliferation

We first determined the Pkm1 and Pkm2 protein levels in proliferating and differentiating C2C12 myoblasts. Pkm1 levels increase significantly as differentiation proceeds (**Fig. 1A**). Pkm2 protein levels also increased at the onset of differentiation, peaking at 24h post-induction of differentiation, but then decreased as differentiation progressed and completed by 72 h (**Fig. 1A**). These results are largely consistent with prior studies in tissue and in cultured cells (33–37).

**Figure 1.**
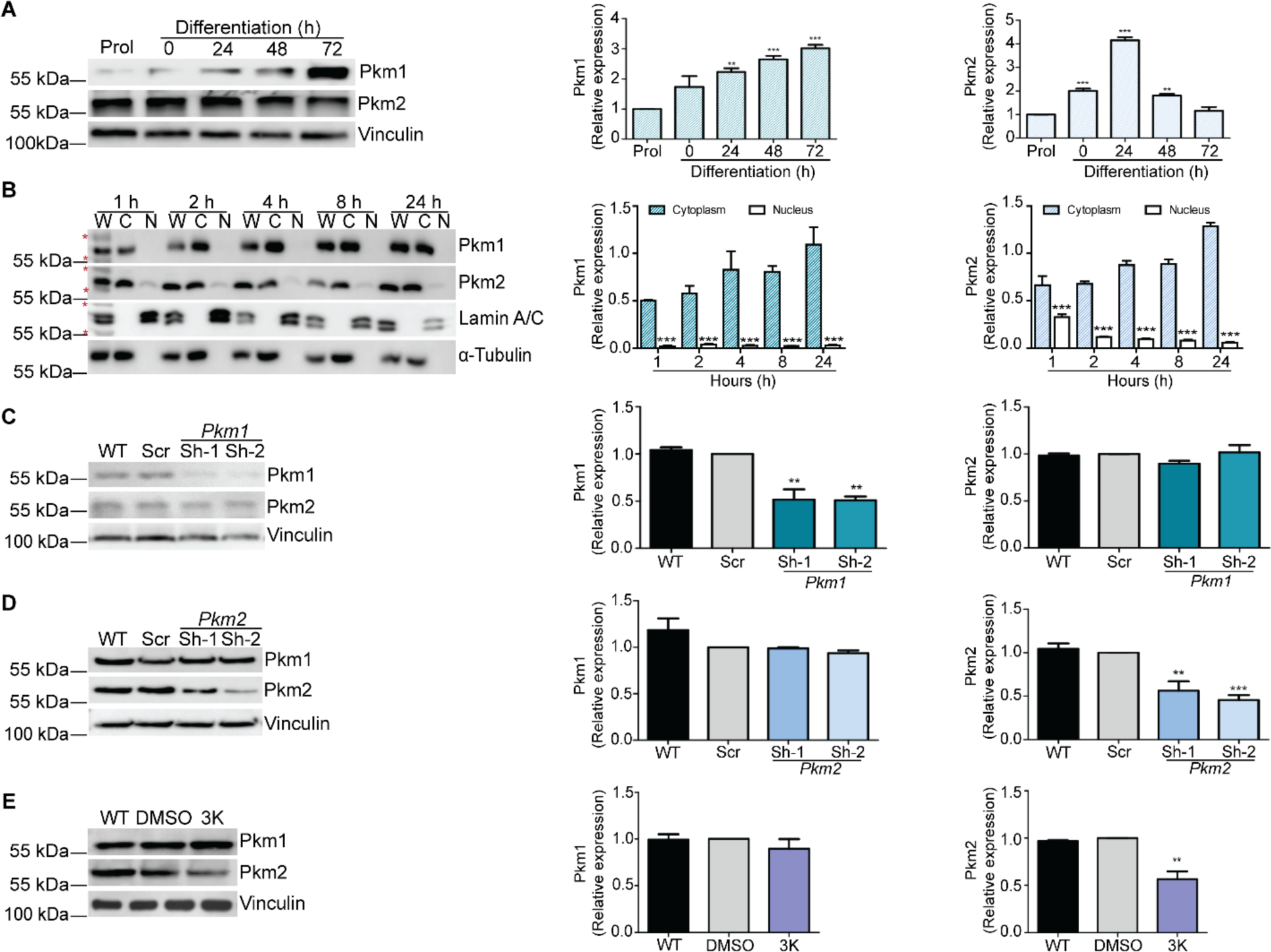
Expression of Pkm1 and Pkm2 during myoblast differentiation. **(A)** Representative western blots (left) and quantification of three independent experiments (right) show Pkm1 and Pkm2 expression in proliferating and differentiating myoblasts. Western blots against vinculin were used as loading controls. **(B)** Subcellular fractionation of differentiating myoblasts was performed to compare Pkm1 and Pkm2 levels in whole cell extract (W) to cytosolic (C) and nuclear (N) fractions. Representative immunoblots (left) and quantification (right) are shown. Immunoblots against Lamin A/C and α-Tubulin were used as controls to show the purity of the fractions. Red asterisks represent molecular-weight size markers. (**C-E**) Representative Western blots (left) and quantification (right) of Pkm1 and Pkm2 levels in *Pkm1* **(C)**, and *Pkm2* **(D)** knockdown samples and 3K inhibitor treated samples **(E)**. Western blots against vinculin were used as loading controls. The samples were compared with the corresponding scrambled and DMSO samples, respectively. For all samples, data are the mean ± SE of three independent biological replicates. **P < 0.01; ***P < 0.001. WT, wild type, Scr, scrambled sequence shRNA.

Pkm1 and Pkm2 are cytoplasmic but have been reported to be nuclear in some cell types (42, 60, 61). We fractionated differentiating myoblasts into nuclear and cytoplasmic fractions and determined the presence of Pkm1 and Pkm2 in each fraction by western blot. We observed clear evidence of Pkm2 in the nuclear fractions, but while the quantification of Pkm1 was above background, we were unable to visualize a Pkm1 band in the nuclear fractions (**Fig. 1B**). We conclude that a fraction of Pkm2 is nuclear in differentiating myoblasts but did not determine if there was no nuclear Pkm1 or if nuclear Pkm1 exists but was below the level of detection in our assay.

To explore the roles of Pkm1 and Pkm2 in myoblasts, we created shRNA vectors to reduce the levels of the Pkm1 or Pkm2 proteins. Knockdown vectors were specific for the target proteins as shown by the western blots and quantification in **Figs. 1C-D**. We supplemented our studies with a Pkm2-specific chemical inhibitor, the 3K compound, that locks Pkm2 into a conformation associated with low activity (53). We determined that the appropriate working concentration of 3K was 2 µM, as this represents a sub-lethal dose for myoblasts as defined by a classic trypan blue cell viability assay (**Supp. Fig. 1A**). This dose also affected myoblast differentiation, as shown by a decrease in the formation in myotubes (**Supp. Fig. 1B**). Treatment of myoblasts with 2 μM of the 3K inhibitor reduced the protein expression of Pkm2, with no effect on Pkm1 protein levels (**Fig. 1E ; Supp. Fig. 1C-D**). Western blot analyses further determined that 2 µM 3K did not induce activated caspase 3, a marker of apoptosis (**Supp. Fig. 1C**). Thus the 3K compound affects both Pkm2 function as well as its expression in these cells.

To determine whether Pkm1 or Pkm2 affected myoblast proliferation, we knocked each down in myoblasts and performed cell counts. *Pkm2* knockdown reduced cell proliferation, while *Pkm1* knockdown had no effect (**Fig. 2A**). The 3K inhibitor of Pkm2 reduced myoblast proliferation to a similar extent as the *Pkm2* knockdown (**Fig. 2A**). Prior studies in cancer and non-transformed cells showed that Pkm2 has an essential role in cell cycle control through the transcriptional regulation of cell-cycle-specific genes (43, 62–64). Analysis of asynchronous proliferating C2C12 myoblasts knocked down for *Pkm2* showed no difference in the distribution of cells in different stages of the cell cycle compared to the control cells and a very slight but statistically significant redistribution in the presence of the 3K inhibitor (**Fig. 2B**). *Pkm2* knockdown myoblasts and myoblasts treated with the 3K inhibitor subsequently were subjected to mitotic arrest with the microtubule inhibitor nocodazole; all cells arrested appropriately as shown by propidium iodide staining (**Fig. 2C – left panel**). However, *Pkm2* knockdown myoblasts and myoblasts where the kinase activity was inhibited with 3K were unable to exit mitosis 24 h after removal from the nocodazole (**Fig. 2C -right panel**), suggesting a specific role for Pkm2 in re-entry to the cell cycle after a nocodazole block.

**Figure 2.**
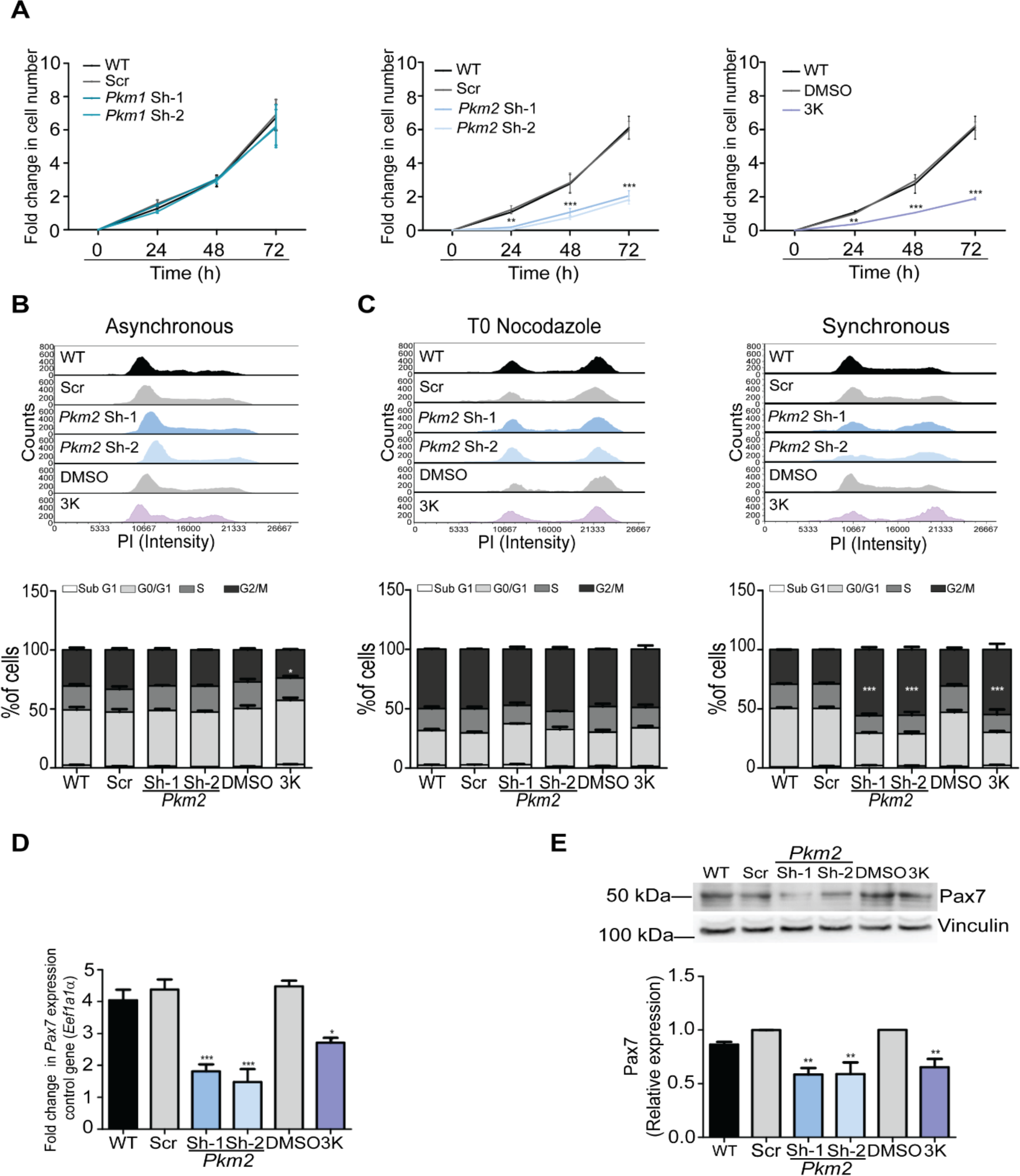
*Pkm2* KD impairs myoblast proliferation by dysregulating cell cycle progression and Pax7. **(A)** Cell counting assay of proliferating wild type (WT) myoblasts, Scr (scrambled shrRNA), *Pkm2* or *Pkm1* knockdown or 3K inhibitor-treated samples. **(B)** Representative histograms of cell cycle progression and the percentage of cells in each cell cycle phase for asynchronously proliferating myoblasts. (**C**) Representative histograms of cell cycle progression and the percentage of cells in each cell cycle phase in cells arrested in mitosis with nocodazole at the time of release (T0), and 24 h post-release. **(D)** Steady state mRNA levels of *Pax7* in proliferating C2C12 myoblasts determined by qRT-PCR. **(E)** Representative western blots (top) and quantification (bottom) showing Pax7 expression in *Pkm2* KD myoblasts and in myoblasts grown in the presence of the 3K inhibitor. For all samples, data are the mean ± SE of three independent biological replicates. *P < 0.05; **P < 0.01; ***P < 0.001. WT, wild type; Scr; scrambled sequence shRNA.

### Pkm2 regulates the expression of the myoblast proliferation marker, Pax7

Pax7 is required for the specification of satellite cells, and for the maintenance, survival, and proliferation of myoblasts (65–67). To determine if the reduction in proliferation observed in myoblasts expressing *Pkm2* shRNA or treated with the 3K inhibitor of Pkm2 was due to an effect on Pax7, we evaluated the expression of *Pax7* mRNA and Pax7 protein. qRT-PCR (**Fig. 2D)**, western blot (**Fig. 2E**) and immunohistochemistry (IHC) analysis (**Supp. Fig. 2**) showed that the steady state mRNA and protein expression levels were reduced by inhibition of *Pkm2* expression and Pkm2 function compared to the control myoblasts.

### Pkm1 and Pkm2 are required for myoblast differentiation

Myoblasts expressing shRNAs that target *Pkm1* or *Pkm2* or that were treated with the Pkm2 inhibitor, 3K, were subjected to standard in vitro differentiation protocols. Images stained at 48 h post-differentiation with the early myogenic marker, myogenin, showed that each of these experimental manipulations had only a modest effect on myogenin staining (**Fig. 3A**). Immunostaining with myosin heavy chain (MHC) showed that each experimental manipulation blocked the formation of multi-nucleated myotubes, indicating non-redundant roles for Pkm1 and Pkm2 in myoblast differentiation in culture. However, *Pkm1* knockdown had a more significant effect on MHC expression than did *Pkm2* knockdown or Pkm2 inhibition (**Fig. 3B**). The inhibition of myotube formation was confirmed and quantified by evaluating myoblast fusion (**Fig. 3C**).

**Figure 3.**
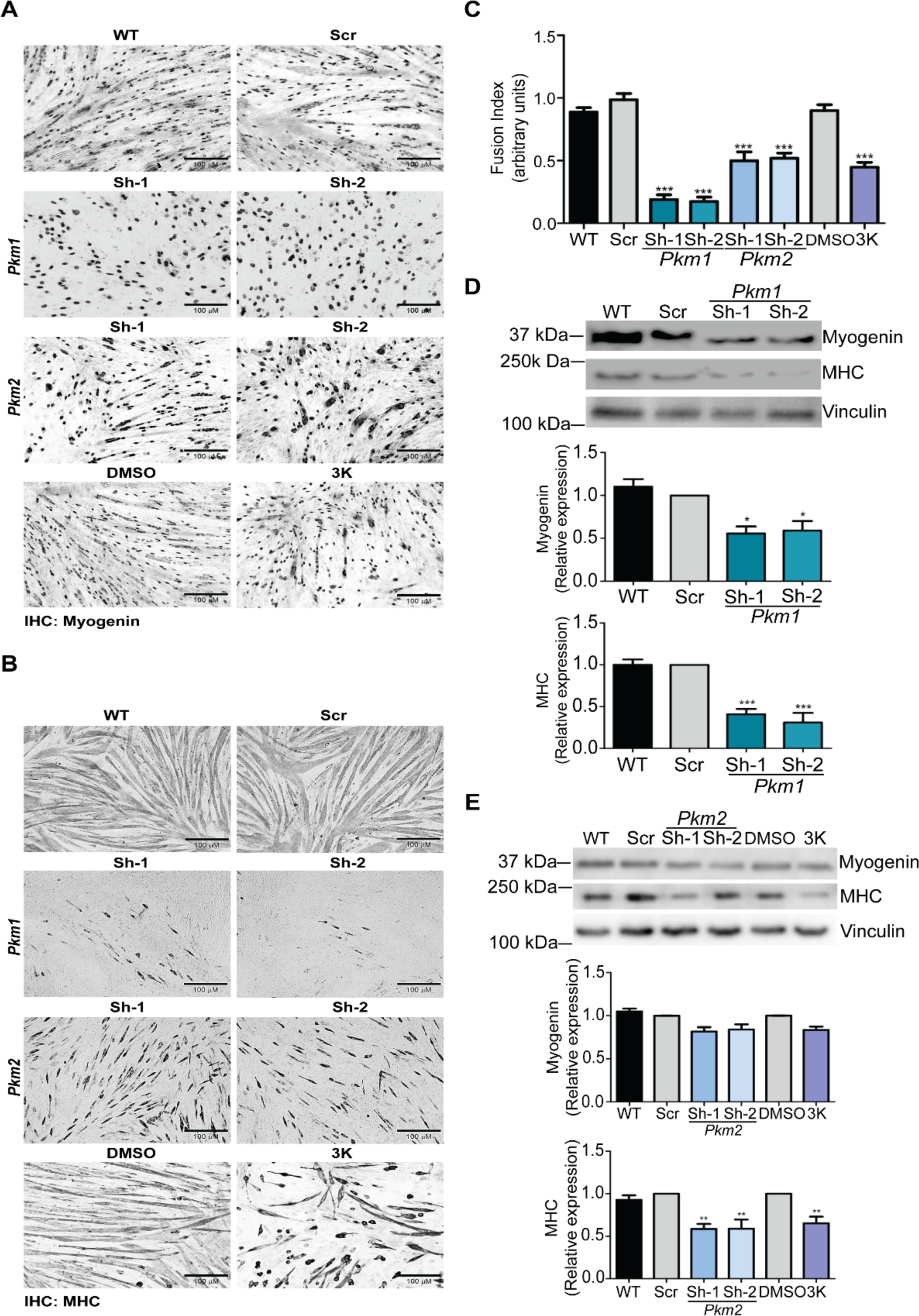
*Pkm1 or Pkm2* KD or Pkm2 inhibition impairs myoblast differentiation. Representative light micrographs of 48 h differentiating myoblasts (WT, Scr, *Pkm1* or *Pkm2* KD or cultured with the 3k inhibitor), immunostained against (**A**) myogenin or (**B**) MHC. (**C**) The fusion index was calculated from the MHC stained samples. (**D-E**) Representative immunoblots and quantification of myogenin and MHC levels in *Pkm1* knockdown samples (**D**), and *Pkm2* knockdown or Pkm2 inhibited samples (**E**). For all samples, data are the mean ± SE of three independent biological replicates. *P < 0.05; **P < 0.01; ***P < 0.001. WT, wild-type; Scr; scrambled sequence shRNA.

Myogenin and MHC protein levels were examined by western blot (**Figs. 3D-E, left panels**) and quantified (**Figs. 3D-E, right panels**). The steady state levels of both myogenin and MHC were reduced by *Pkm1* knockdown, but *Pkm2* knockdown or treatment with 3K (**Fig. 3C**) only affected MHC protein levels, not myogenin protein levels, possibly suggesting some specificity by Pkm2 in the regulation of protein levels from different genes. When steady state mRNA levels were examined by qPCR, the expression of both *myogenin* and an MHC isoform (*MyhIIb*), as well as four other genes that are induced or elevated during myoblast differentiation, were reduced by *Pkm1* or *Pkm2* knockdown or by 3K treatment (**Supp. Fig. 3**). The data suggest that transcription of myogenic genes requires both Pkm1 and Pkm2 and indicate that the Pkm isoforms do not have redundant functions in myogenic gene expression during differentiation in culture.

### Distinct transcriptional effects of *Pkm1* and *Pkm2* knockdown during myoblast differentiation

Our current data and prior evidence show differential temporal expression of Pkm1 and Pkm2 *in vitro* and *in vivo* (33–37), suggesting the possibility of differential impacts on gene expression. We used RNA-seq to investigate global changes in gene expression patterns in differentiating myoblasts that result from knockdown of each isoform. Reads were mapped to the murine genome (mm10) and gene expression levels were determined. Read information and correlation coefficient of replicates is presented in **Supp. Table 3**; the Pearson correlation coefficient for the replicate samples of sh*Scr*, sh*Pkm1*, and sh*Pkm2* were 0.986 or greater. Significant differential gene expression in both replicates for each shRNA was defined as log2FoldChange >1 and <−1, padj < 0.05. **Supp. Table 4** shows the differentially expressed genes for each shRNA.

518 genes were differentially regulated upon *Pkm1* knockdown while 952 genes were differentially regulated by *Pkm2* knockdown. A significant number of genes (318) were targets of both knockdowns, leaving 200 genes uniquely affected by *Pkm1* knockdown and 634 genes uniquely affected by *Pkm2* knockdown (**Fig. 4A**). We performed GO (Gene Ontology) analysis of differentially expressed genes uniquely affected by *Pkm1* or *Pkm2* knockdown as well as differentially expressed genes that were regulated by either *Pkm1* or *Pkm2* knockdown (**Supp. Table 5**). The top ten GO terms for genes that were down-regulated by *Pkm2* knockdown were all related to skeletal muscle differentiation or function, while seven of the top ten GO terms for genes down-regulated by either *Pkm1* or *Pkm2* knockdown were related to skeletal muscle differentiation or function (**Fig. 4B**). In contrast, genes uniquely down-regulated by *Pkm1* knockdown were heavily weighted toward terms that are related to the metabolic functions of the Pkm enzymes, namely ATP synthesis, ATP-dependent processes integral to cell viability, and functions related to respiration (**Fig. 4B**). GO analysis of genes that were up-regulated by *Pkm1* knockdown or by either *Pkm1* or *Pkm2* knockdown identified terms related to extra-cellular matrix, adhesion, and cytoplasmic filament organization. Analysis of genes uniquely induced by *Pkm2* knockdown identified terms mostly related to cell division (**Fig. 4C**), consistent with the observations that Pkm2 primarily impacted myoblast proliferation (**Fig. 2**).

**Figure 4.**
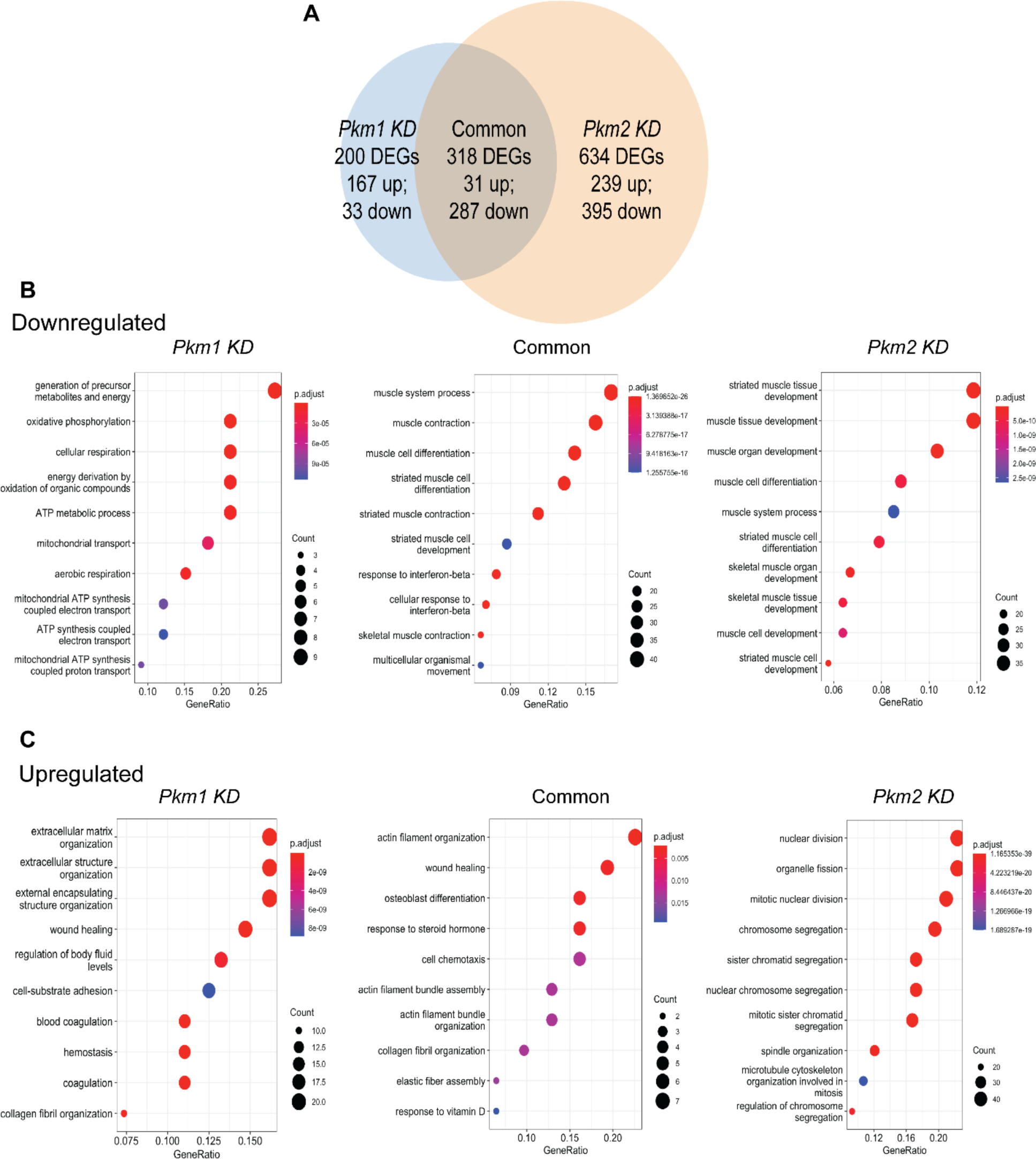
Differential gene expression resulting from *Pkm1* or *Pkm2* KD during myoblast differentiation. **(A)** Venn diagram showing the unique and overlapping differentially expressed genes (DEGs) between differentiating myoblasts KD for *Pkm1* and *Pkm2* using Scr shRNA samples as control. GO term analysis for downregulated **(B)** and upregulated **(C)** genes. Analysis of differentially expressed genes in differentiating C2C12 cells *Pkm1* KD (left panel) or *Pkm2* KD (right panel) or overlapping common genes (central panel). Cut-off was set at 2.0 of the −log (adjusted p value). See Supp. Table 4 for the complete list of genes.

The results indicate overlap between the roles of Pkm1 and Pkm2 in myoblast differentiation but also that each Pkm has some specificity. Of note, Pkm1 uniquely regulated genes involved in metabolism while Pkm2 uniquely regulated some genes involved in differentiation. This likely explains the requirement for both Pkm1 and Pkm2 in the differentiation of C2C12 myoblasts.

### Pkm2 specifically regulates the expression of mSWI/SNF chromatin remodeling enzyme subunits belonging to the cBAF subfamily that is required for myoblast differentiation

We noted that among the genes specifically down-regulated by *Pkm2* KD was the *Dpf2* gene, which encodes the mSWI/SNF chromatin remodeling enzyme subunit called Dpf2/Baf45d. The family of mSWI/SNF chromatin remodeling enzymes can be classified into three major sub-families called consensus BAF (cBAF), non-consensus BAF (ncBAF), and polybromo-containing BAF (PBAF) (50). The three sub-families have shared as well as unique subunits, and each of the sub-families have been linked to specific functions (68–72). The finding that *Dpf2* expression is down-regulated by *Pkm2* knockdown is of interest because the Dpf2 subunit is unique to the cBAF sub-family, which we have previously shown is responsible for skeletal muscle-specific gene expression during myoblast differentiation (51). This suggests a mechanism for the observed effects of *Pkm2* knockdown on myoblast differentiation.

We confirmed that *Dpf2* expression was specifically downregulated by *Pkm2* knockdown cells by qPCR (**Fig. 5A**). We also examined expression of the gene that encodes the gene expressing Baf250a, another subunit unique to the cBAF sub-family of complexes (50, 73, 74), along with the expression of the gene that encodes the Brg1 ATPase subunit that is found in all three mSWI/SNF sub-families (50, 75–77) and that is required for myogenic differentiation (11). Like *Dpf2*, the gene encoding Baf250A was down-regulated specifically in *Pkm2* knockdown cells (**Fig. 5B**), whereas the gene encoding the Brg1 ATPase subunit was not affected by either *Pkm1* or *Pkm2* knockdown (**Fig. 5C**). We subsequently evaluated protein expression. *Pkm1* knockdown did not affect the expression levels of Dpf2, Baf250a or Brg1 proteins (**Fig. 5D**). *Pkm2* knockdown reduced the protein levels of Dpf2 and Baf250a, (**Fig. 5E**), consistent with the gene expression data. Although *Pkm2* knockdown did not impact Brg1 mRNA expression, Brg1 protein levels were reduced. Inhibition of Pkm2 function using the 3K inhibitor gave the same results as observed upon *Pkm2* knockdown. *Dpf2* and the gene encoding Baf250a were down-regulated in cells in which Pkm2 was inhibited, as were Dpf2 and Baf250a protein levels (**Supp. Fig. 4**). Brg1 mRNA was unaffected by Pkm2 inhibition, but Brg1 protein levels were reduced (**Supp. Fig. 4**). These observations indicate that Pkm2 can impact the expression of multiple mSWISNF subunits involved in myogenic differentiation and can do so by affecting mRNA and protein accumulation.

**Figure 5.**
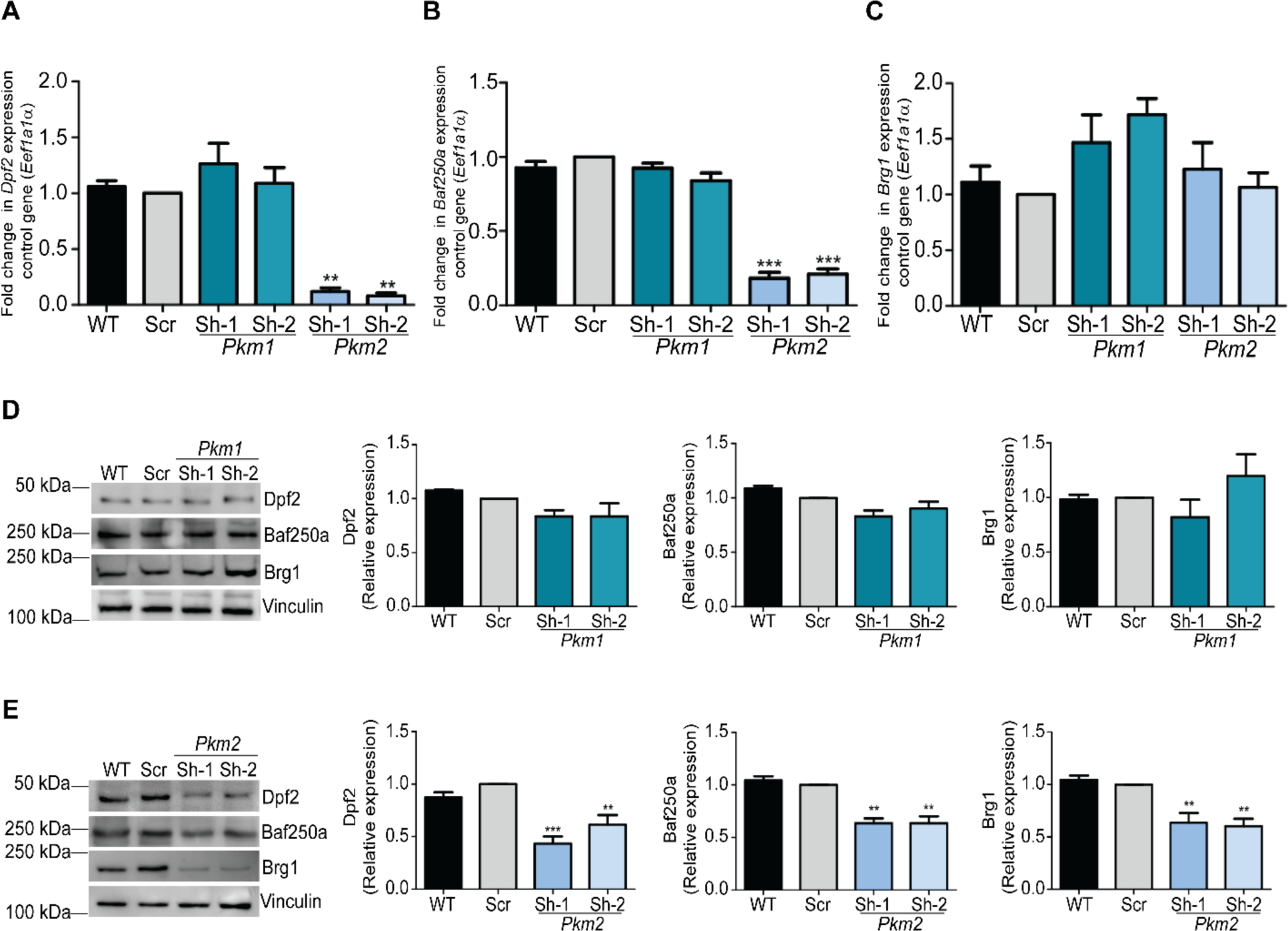
*Pkm2* KD downregulates Baf250a and Dpf2 expression in differentiating C2C12 cells. Steady state mRNA levels of **(A)** *Dpf2*, **(B)** *Baf250a*, and **(C)** *Brg1*, determined by qRT-PCR from 48 h differentiating C2C12 myoblasts. **(D)** Representative immunoblot (left) and quantification (right) of Dpf2, Baf250a, and Brg1 protein levels in differentiating control cells and *Pkm1* KD myoblasts. Immunoblots against vinculin were used as loading controls. **(E)** As in (D) except *Pkm2* KD cells were analyzed, 48 h after inducing myoblast differentiation. The data represent three independent biological experiments. Bar graphs show the mean ± SE. *P < 0.05, **P < 0.01, and ***P < 0.001. WT, wild-type; Scr; scrambled sequence shRNA.

### Pkm1, but not Pkm2, affects the sub-cellular localization of Dpf2

The mSWI/SNF chromatin remodeling enzymes are nuclear proteins. However, when differentiated myoblasts were separated into nuclear and cytoplasmic fractions, we observed that even though *Pkm1* knockdown did not affect *Dpf2* mRNA or Dpf2 protein levels, a significant portion of the Dpf2 protein in these cells was present in the cytoplasm (**Fig. 6A**). Moreover, the migration of the cytoplasmic Dpf2 protein in the gel was slightly greater than what was observed for the nuclear Dpf2 protein in the *Pkm1* knockdown and the control cells (**Fig. 6A**). This observation correlates with the reduced level of Pkm1 and suggests that Pkm1 phosphorylation of Dpf2 may be needed for nuclear localization. We also examined Baf250a and Brg1 in the nuclear and cytoplasmic fractions. *Pkm1* knockdown had no effect on the nuclear localization of these mSWI/SNF subunits; all of the Baf250a and Brg1 was located in the nuclear fractions. Quantification of the results are presented in the graphs shown in **Fig. 6A**.

**Figure 6.**
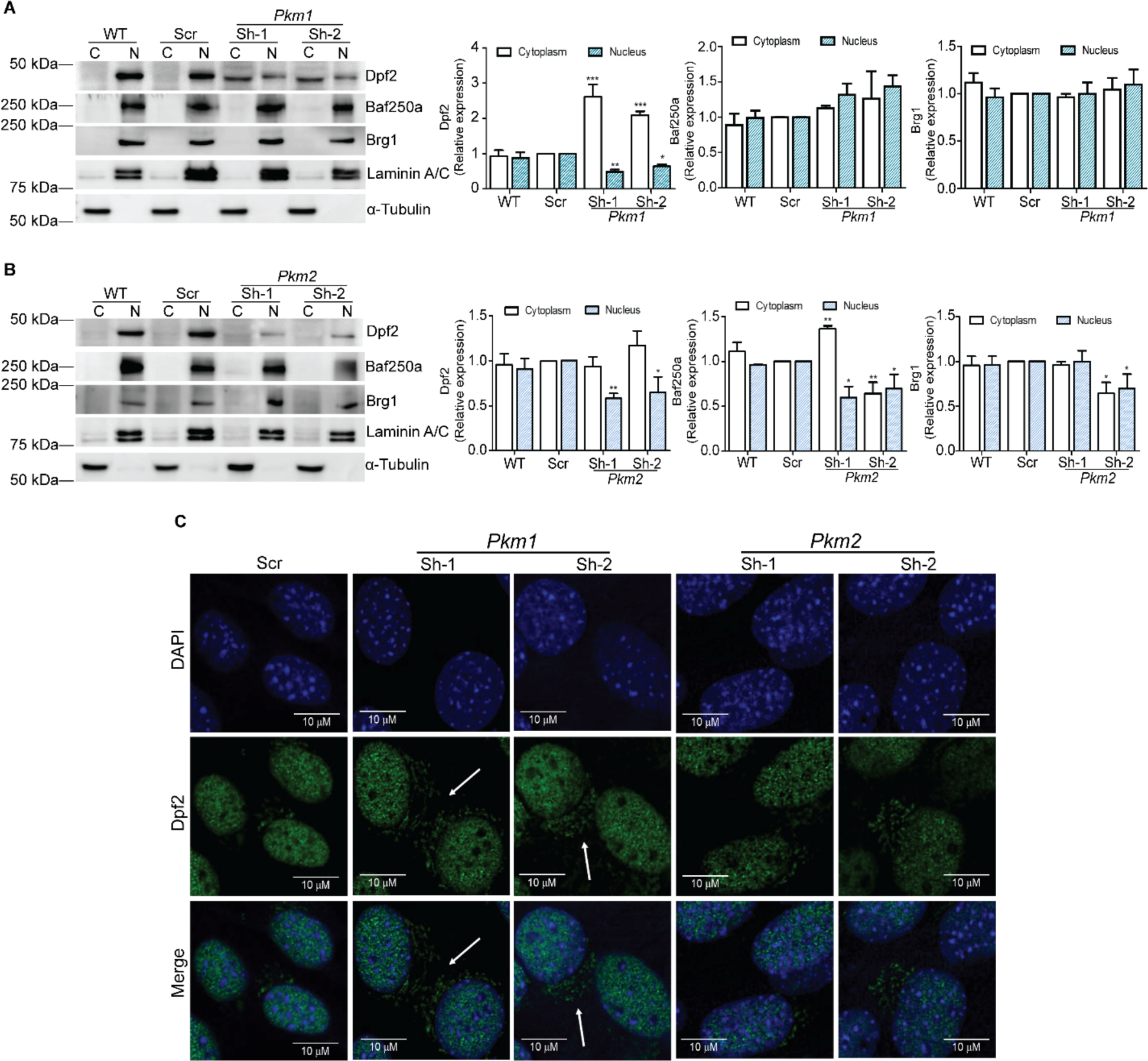
Pkm1 regulates the sub-cellular localization of Dpf2 in differentiating C2C12 cells. Subcellular fractionation of myoblasts differentiated for 48h was performed to compare cytosolic (C), and nuclear (N) fractions. **(A-B)** Representative immunoblots (left) and quantification (right) of Dpf2, Baf250a, and Brg1 levels in WT and Scr controls as well as in Pkm1 KD **(A)**, and Pkm2 KD cells **(B)**. Immunoblots against Laminin A and α-Tubulin were used as controls to show the purity of the fractions. The data represent three independent biological experiments. Bar graphs show the mean ± SE. *P < 0.05, **P < 0.01 or ***P < 0.001. **(C)** Confocal images of differentiated C2C12 myoblasts (48 h) immunostained with a specific anti-rabbit antibody against Dpf2 (green). Nuclei stained with DAPI (blue). Scale bar: 10 μm. WT, wild-type; Scr; scrambled sequence shRNA.

Repetition of this experiment with *Pkm2* knockdown differentiated myoblasts showed no effect on nuclear localization of Dpf2, Baf250a, or Brg1 (**Fig. 6B**). However, the reduction in the amount of each of these three proteins (**Fig. 5, Supp. Fig. 4**) was confirmed. Immunostaining of the differentiated myoblasts revealed extra-nuclear staining in the *Pkm1* knockdown cells and exclusively nuclear staining in the Scr control cells and in the *Pkm2* knockdown cells (**Fig. 6C**). These results confirm those obtained via biochemical approach used in **Figs. 5 and 6A -B**. We conclude that both isoforms of Pkm regulate the Dpf2 protein; Pkm1 contributes to Dpf2 nuclear localization while Pkm2 regulates *Dpf2* expression.

### Pkm2 regulates the binding of Dpf2 and Baf250a at the *Myogenin* and *MyhcIIb* promoters

*Pkm2* knockdown reduced the expression of the Dpf2 and Baf250a protein, while *Pkm1* knockdown reduced the amount of nuclear Dpf2. mSWI/SNF enzymes bind to myogenic promoters when myogenic genes are activated during differentiation. We performed chromatin immunoprecipitation (ChIP) experiments to determine whether *Pkm1* or *Pkm2* knockdown impair the binding of Dpf2 or Baf250a to myogenic promoters controlling the *myogenin* and the *MyhcIIb* genes. *Pkm2* knockdown reduced the binding of both Dpf2 and Baf250a to both promoters (**Fig. 7A-B**). *Pkm1* knockdown resulted in a statistically significant decrease in Baf250a binding at the *MyhcIIb* promoter, but not at the *myogenin* promoter. *Pkm1* knockdown also did not impact Dpf2 binding at either promoter (**Fig. 7A-B**). The data suggest that Pkm2 regulates the ability of these mSWI/SNF subunits to bind to myogenic promoters and that Pkm1 minimally affects mSWI/SNF subunit binding.

**Figure 7.**
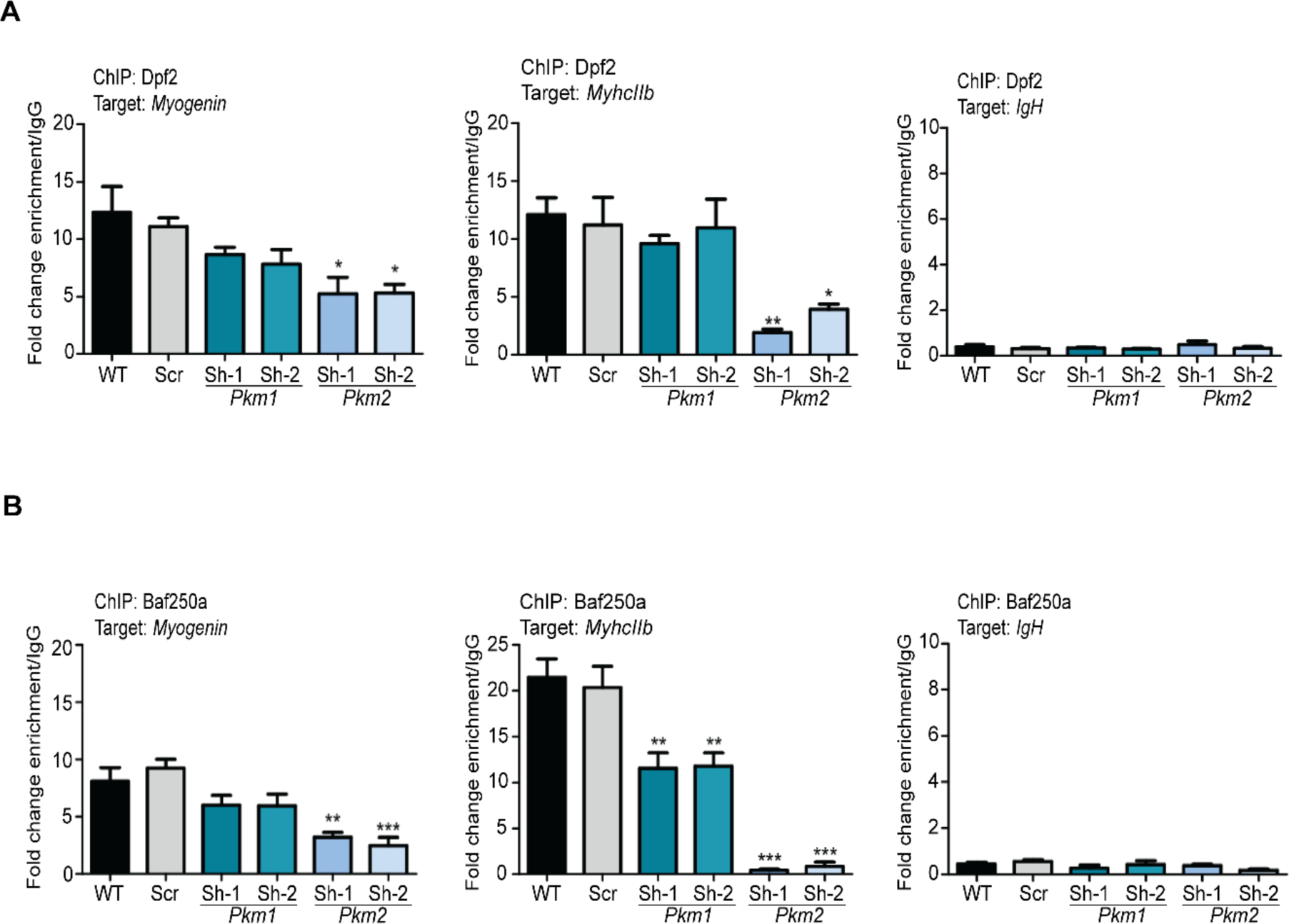
Pkm1 and Pkm2 differentially regulate the binding of Dpf2 and Baf250a at the *Myogenin* and *MyhcIIb* promoters. ChIP-qPCR showing the binding of **(A)** Dpf2 or **(B)** Baf250a to the *Myogenin* and *MyhIIb* promoters in differentiating myoblasts. The *IgH* promoter was used as negative control target for binding. The data was normalized using IgG as a negative control for the ChIP. The data are presented as the mean ± SE from three independent biological experiments. *P < 0.05; **P < 0.01; ***P < 0.001. WT, wild-type; Scr; scrambled sequence shRNA.

### Pkm2 knockdown reduces H3 phosphorylation at T6, T11 and T45

Pkm2, but not Pkm1, was specifically linked to H3 phosphorylation at T11 in work showing H3T11, but not H3T3 or H3T6, was a direct target for Pkm2 in cancer cells (43). H3T11 phosphorylation has not been characterized in myoblast differentiation. The H3 tail domain is also phosphorylated at other T residues, including T45. A time course assessment of the levels of each of the H3 modifications during myoblast differentiation was performed. Phosphorylated H3T6 levels were similar in proliferating and differentiating myoblasts, with a statistically significant increase at 72h, a time when differentiation is largely complete (**Fig. 8A**). Levels of phosphorylated H3T11 and T45, in contrast, had increased at the onset of differentiation relative to the level present in proliferating myoblasts, reached peak levels 24-48 h after the onset of differentiation, and showed a slight decline by 72 h (**Fig. 8A**). Knockdown of *Pkm2* or inhibition of Pkm2 by the 3K compound reduced the level of phosphorylated H3T11, as might be expected, but also decreased the levels of phosphorylated H3T6 and H3T45 (**Fig. 8B**). These results raised the possibility of indirect regulation of these phosphorylation events by Pkm2. Bulk levels of phosphorylated H3T6, H3T11, and H3T45 did not change upon *Pkm1* knockdown **(Fig. 8C)**.

**Figure 8.**
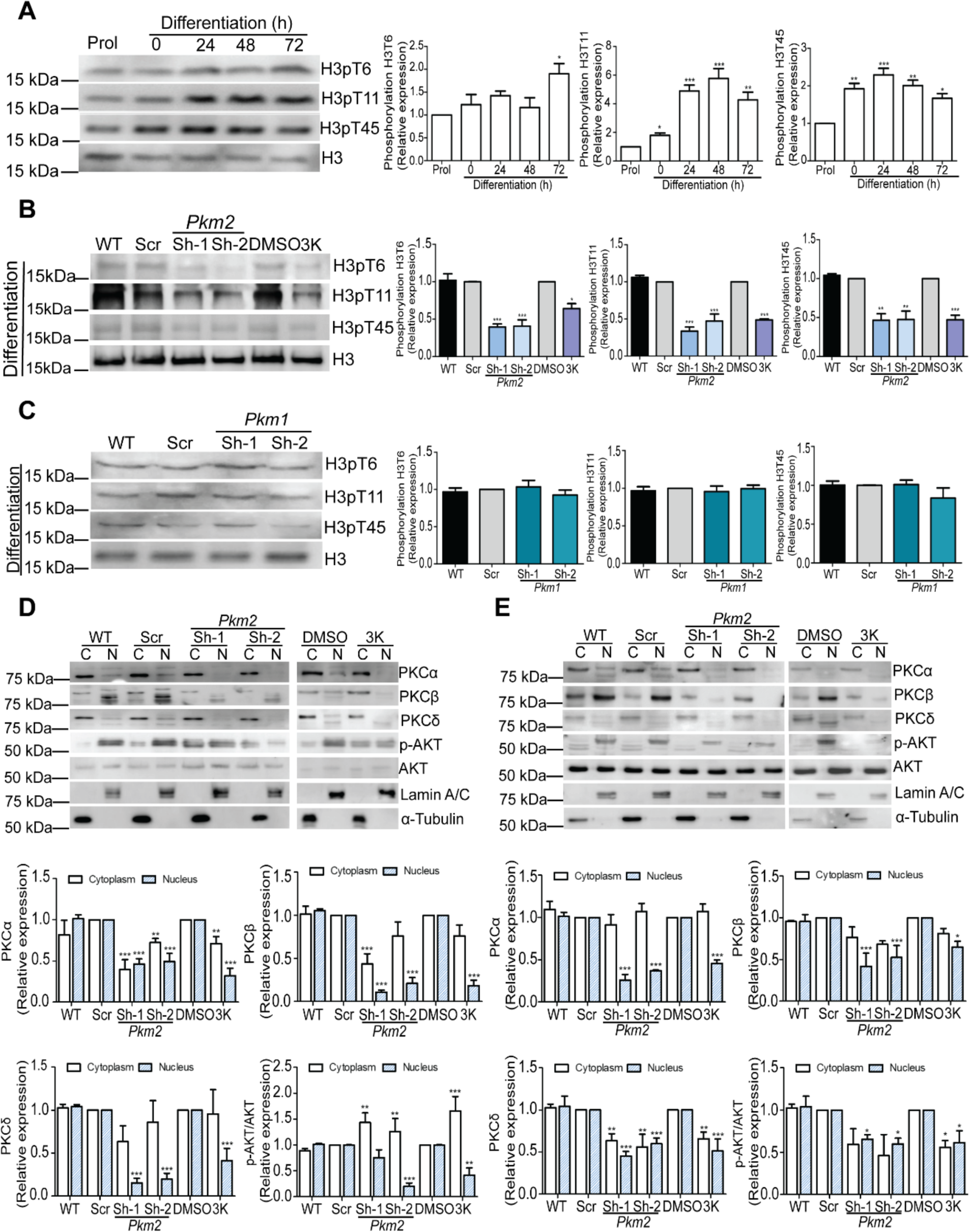
*Pkm2* KD or Pkm2 inhibition decreases both bulk levels of phosphorylated histones H3T6, H3T11, and H3T45 and regulates the kinases that promote the phosphorylation of H3T6 and H3T45. **(A)** Representative western blots (left) and quantification (right) show H3T6, H3T11, and H3T45 phosphorylation in proliferating myoblasts and in myoblasts at different stages of differentiation. **(B-C)** Representative western blots (left) and quantification (right) show H3T6, H3T11, and H3T45 phosphorylation in *Pkm2* KD myoblasts or in the presence of the 3K inhibitor (B) or *Pkm1* KD (C) in 48 h differentiating myoblasts. Immunoblots against total H3 were used as loading controls. **(D-E)** Representative western blots (top) and quantification (bottom) show PKCα, PKCβ, PKCδ and p-ATK expression in nuclear (N) and cytoplasmic (C) fractions in proliferating (D) or differentiating (E) myoblasts upon *Pkm2* KD or in the presence of the 3K inhibitor. Samples were compared to the corresponding Scr or DMSO sample. The data represent three independent biological experiments. Bar graphs show the mean ± SE. *P < 0.05; **P < 0.01; ***P < 0.001. Prol.; proliferating cells, WT, wild-type; Scr; scrambled sequence shRNA.

### Pkm2 regulates downstream kinases that regulate H3T6 and H3T45 phosphorylation

A decrease in the bulk levels of phosphorylated H3T11 in the presence of *Pkm2* knockdown or Pkm2 inhibition was not surprising given prior work demonstrating Pkm2 can phosphorylate H3T11 in malignant glioma cells expressing EGFR (43). However, the same study showed that Pkm2 did not phosphorylate H3T6, suggesting that manipulation of Pkm2 indirectly affects the bulk levels of phosphorylated H3T6 and, possibly, bulk levels of phosphorylated H3T45 as well.

PKCβ enzyme is known to phosphorylate H3T6 (46), and both PKCδ and phospho-AKT can phosphorylate H3T45 (47, 49). We therefore examined levels of PKC isoforms and phospho-AKT in nuclear and cytoplasmic fractions from proliferating **(Fig. 8D)** and differentiating **(Fig. 8E)** myoblasts subjected to *Pkm2* knockdown or Pkm2 inhibition. The amounts of nuclear PKCα, β and δ and phospho-AKT were reduced upon either form of Pkm2 manipulation, regardless of the proliferation or differentiation status of the myoblasts **(Fig. 8D-E)**. The impact of *Pkm2* knockdown or Pkm2 inhibition on cytoplasmic levels of these kinases was more limited. The only consistent results were a decrease in cytoplasmic PKCα levels in proliferating cells and a decrease in cytoplasmic PKCδ levels in differentiating cells **(Fig. 8D-E)**. These findings support the idea that the reduction in the levels and in the chromatin incorporation of phosphorylated H3T6 and H3T45 due to *Pkm2* knockdown or Pkm2 inhibition results from an indirect effect of Pkm2 on the kinases that directly phosphorylate H3T6 and H3T45.

### Pkm2 mediates incorporation of phosphorylated H3T6, H3T11, and H3T45 into myogenic regulatory sequences during myoblast differentiation

Prior evidence indicated that phosphorylated H3T6, T11, and T45 are associated with gene activation and are incorporated at actively expressing genes (46, 48, 49). We performed ChIP experiments that revealed an increase in incorporation of these phosphorylated H3 molecules at the *myogenin* promoter at the onset of differentiation in the case of H3T6 and H3T11, with increased incorporation of H3T45 being slightly delayed, but still occurring in the initial hours of differentiation (**Supp. Fig. 5A**). Similar analyses were performed on four other myogenic regulatory sequences: the *MyhcIIb*, *Cav3*, and *Tnnt3* promoters as well as the *Ckm* enhancer (**Supp. Fig. 5B-E**). While the different regulatory sequences all showed an increase in the incorporation of each of the phosphorylated H3 molecules, the time of the initial increase and whether the increase was sustained or transient varied with each phosphorylated H3 species and by gene regulatory sequence **(Fig. 9B-E)**.

**Figure 9.**
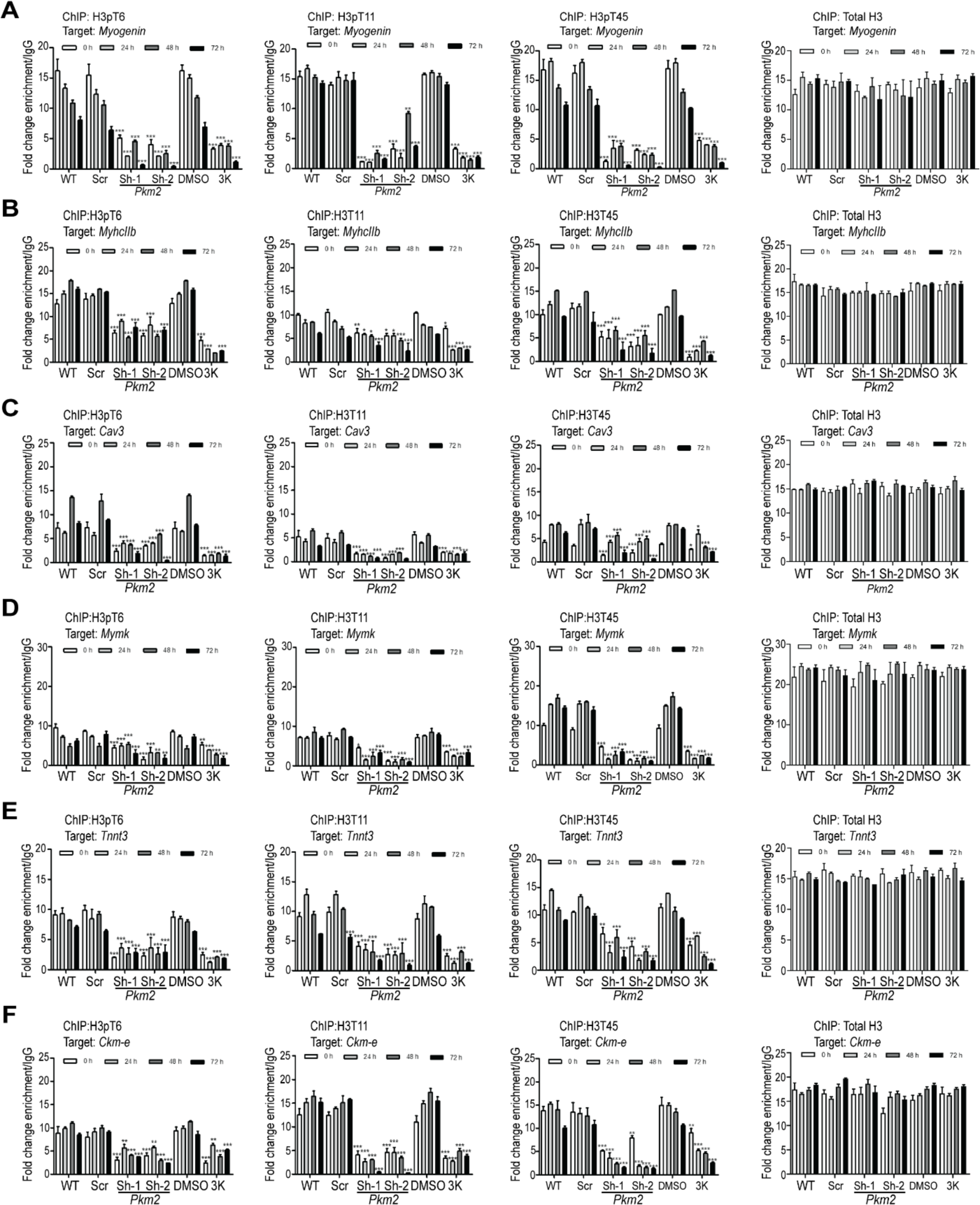
*Pkm2* KD and Pkm2 inhibition decrease the incorporation of phosphorylated histones H3T6, H3T11, and H3T45 at myogenic promoters. ChIP-qPCR showing binding of phosphorylated H3T6, H3T11, H3T45 or total H3 to the (A) *Myogenin*, (B) *MyHCIIb,* (C) *Cav3*, (D) *Mymk, and* (E) *Tnnt3* promoters and **(F)** the *Ckm* enhancer (*Ckm-e*) in differentiating myoblasts at the indicated times. The ChIP data were normalized using IgG as a control for the ChIP. The data for all experiments represent the mean ± SE of three independent biological experiments. *P < 0.05; **P < 0.01; ***P < 0.001. WT, wild type, Scr, scrambled shRNA.

Assessment of the incorporation of these specifically phosphorylated H3 molecules at multiple myogenic regulatory sequences in myoblasts differentiated in the presence of *Pkm2* knockdown or in the presence of the Pkm2 inhibitor was performed. The five regulatory sequences examined in Supp. Fig. 5 and the *Myomaker* (*Mymk*) promoter were analyzed. Incorporation of phosphorylated H3T6, H3T11, and H3T45 was reduced at each of the tested myogenic regulatory sequences across the 72 h time course of differentiation **(Fig. 9A-F)**. The decreased incorporation of phosphorylated H3T11 at the myogenic regulatory sequences is consistent with Pkm2 directly phosphorylating H3T11. The reduction of incorporation of phosphorylated H3T6 and H3T11 at myogenic promoters is consistent with the observed reduction in bulk levels of these two H3 species **(Fig. 8B)**, and further supports the idea that Pkm2 is indirectly affecting phosphorylation of H3T6 and H3T45.

### Pkm2-mediated regulation of phosphorylated H3-T6, T11, and T45 also occurs at the *Pax7* promoter in proliferating cells; regulation is not differentiation-specific

We examined the impact of *Pkm1* or *Pkm2* knockdown and Pkm2 inhibition on the levels of the three phosphorylated H3 species in proliferating myoblasts. *Pkm2* knockdown or Pkm2 inhibition, but not *Pkm1* knockdown, reduced the bulk levels of intra-cellular phosphorylated H3T6, H3T11, and H3T45 **(Supp. Fig. 6A-B)**. Expression of the *Pax7* gene is necessary to maintain the proliferative state in myoblasts (65–67). ChIP experiments demonstrated that all three phosphorylated H3 species are incorporated into the *Pax7* promoter in proliferating cells **(Supp. Fig. 6C)**. Assessment of phosphorylated H3 proteins at the *Pax7* promoter as myoblasts progressed through differentiation showed a progressive decrease in incorporation (**Supp. Fig. 6C)** that correlates with the shutdown of *Pax7* gene expression that is an essential event during differentiation (65). These results further support the correlation between the presence of these phosphorylated forms of H3 at gene regulatory sequences and active gene expression. *Pkm2* knockdown or Pkm2 inhibition showed that incorporation of phosphorylated H3-T6, -T11, and-T45 at the *Pax7* promoter was reduced relative to the controls (**Supp. Fig. 6D)**. The role of Pkm2 in regulating phosphorylated H3T6, H3T11, and H3T45 in proliferating and differentiating myoblasts is therefore similar.

## DISCUSSION

We report novel functions for the muscle-specific isoforms of pyruvate kinase, Pkm1 and Pkm2. Gene expression analyses of differentiating myoblasts revealed that genes uniquely affected by *Pkm1* knockdown were predominantly associated with cell metabolism, consistent with pyruvate kinase’s conventional role in glycolysis. Genes targeted by either Pkm1 or Pkm2 or uniquely targeted by Pkm2 also impacted metabolism, but the top ten GO terms indicated the impacted genes were associated with skeletal muscle differentiation and function. *Pkm2* knockdown or inhibition of the Pkm2 kinase decreased the expression of proteins that are specific for the cBAF subfamily of the mSWI/SNF chromatin remodeling enzymes that we previously showed are required for myoblast proliferation and differentiation (51, 59), suggesting that Pkm2 impacted myogenic gene expression via effects on the chromatin remodeling enzyme important for differentiation-specific gene expression (51). By contrast, *Pkm1* knockdown did not affect cBAF subunit expression. However, it decreased the nuclear:cytoplasmic ratio of the Dpf2 subunit, lowering the nuclear concentration of this subunit. In addition, Pkm2 was previously identified as a kinase for H3T11, creating a mark generally associated with active transcription (46). We also demonstrated that phosphorylation levels of H3T6 and H3T45 were diminished at myogenic promoters during myoblast proliferation and differentiation upon *Pkm2* knockdown. This suggests an indirect role for Pkm2 in modulating the phosphorylation status of H3T6 and H3T45, potentially by reducing the amount and/or activity of the PKC and AKT kinases that phosphorylate these specific sites on H3 tails. Together, the results provide novel mechanisms for Pkm activity and illuminate the complexities of Pkm function outside of its traditional glycolytic functions.

Despite the apparent plethora of Pkm2-specific functions, is it important to remember that Pkm1 and Pkm2 differ by only the inclusion of one of two 45 amino acid exons (30). It is therefore not surprising, and likely expected, that there should be some redundancy between the two proteins. Most of the down-regulated genes in our RNA-seq analysis that were associated with skeletal muscle development and function were sensitive to either *Pkm1* or *Pkm2* knockdown, suggesting that either enzyme can mediate activation of these genes, at least in the context of cultured myoblasts. Our work is consistent with earlier studies showing primary myoblasts isolated from a conditional mouse knockout of *Pkm2* were unable to differentiate into myotubes in culture when *Pkm2* was depleted prior to differentiation (62). However, whole-body Pkm2 knockout mice were viable and fertile (78), and mice with muscle-stem cell-specific knockout of *Pkm2* showed no defect in skeletal muscle development or regeneration after injury (33). These animal studies suggest a redundancy between Pkm1 and Pkm2 in embryonic and adult myogenesis. The difference between the *in vitro* and *in vivo* results may be due to nutrient availability differences in the tissue culture environment, as has been proposed previously (33). Nevertheless, the existence of the two isoforms and the evidence of Pkm2 functions outside of glycolysis point to the likelihood of important, Pkm2-specific functions.

We determined that the expression of the genes encoding the mSWI/SNF subunits, Dpf2 and Baf250a, were downregulated upon *Pkm2* knockdown. Expression of the gene encoding the ubiquitous ATPase subunit Brg1, however, was not affected by *Pkm2* knockdown. A regulatory role for Pkm2 in the expression of specific subunits of the mSWI/SNF chromatin remodeling enzymes that are required for myoblast proliferation and differentiation was not expected. Yet these finding are entirely consistent with prior findings. Dpf2 and Baf250a are subunits of the cBAF subfamily of the mSWI/SNF chromatin remodeling enzymes but are not present in the other two major sub-families of mSWI/SNF enzymes (50). Our previous studies showed that cBAF enzymes were specifically required for myoblast proliferation and differentiation via regulation of the *Pax7* gene and differentiation-specific gene expression (51, 59). Thus the phenotypes observed due to partial depletion of cBAF-specific subunits upon *Pkm2* knockdown mimicked those identified upon the direct knockdown of the genes encoding the Dpf2 and Baf250a proteins (51, 59). The mechanism(s) of Pkm2 action on cBAF-specific subunit expression remain to be determined. Likely possibilities are Pkm2-mediated phosphorylation of one or more components of the transcriptional machinery that drives gene expression and/or the phosphorylation of histones and/or other non-histone chromatin proteins at the promoters controlling the expression of the cBAF-specific subunits. The latter possibility, however, would likely require a promoter-specific targeting mechanism. Little is known about the regulation of these genes, so this hypothesis remains unexplored.

The mSWI/SNF subunits are phosphoproteins (79–82). To date, p38, AKT, CK2, and PKCβ have been identified as kinases involved in mSWI/SNF subunit phosphorylation and function in myoblast proliferation or differentiation, and calcineurin has been identified as a critical phosphatase that counteracts PKCβ and facilitates the activation of the Brg1 ATPase for chromatin remodeling and myogenic gene induction(15, 24–28, 83). We add Pkm1 and Pkm2 to the list of signaling molecules that can regulate the mSWI/SNF chromatin remodeling enzymes, further increasing the complexity of mSWI/SNF protein modifications.

Pkm2 is known to directly phosphorylate H3T11 in malignant glioma cells expressing EGFR (43). *Pkm2* knockdown reduced phosphorylated H3T11 levels in bulk as well as at myogenic gene regulatory sequences in myoblasts, providing an additional putative mechanism for Pkm2 function during myogenic gene activation. We simultaneously examined the bulk levels of phosphorylated H3T6 and phosphorylated H3T45 as well as the levels of incorporation of these modified histones at myogenic gene regulatory sequences. We anticipated that one or both of these modified histones would be unaffected by *Pkm1* or *Pkm2* knockdown. Instead, we observed decreases in bulk and promoter-specific phosphorylated H3T6 and H3T45, and we attribute these findings to an indirect effect of Pkm2 knockdown on the PKC and AKT kinases that have been implicated as mediating these modifications. A simple explanation is that Pkm2 directly phosphorylates PKC and AKT, but Pkm2-mediated modification of other kinases/phosphatases in the pathways leading to PKC and AKT activity is also possible. Alternately, or in addition, Pkm2 could be phosphorylating other cellular proteins involved in driving the expression or stability of PKC proteins. Prior work determined that Pkm2 did not directly phosphorylate H3T6 (43), but to our knowledge H3T45 has not been tested. Thus another possibility is that Pkm2 directly phosphorylates H3T45.

The primary novel contribution of Pkm1 to myoblast differentiation was its requirement for the nuclear localization of Dpf2, a specific subunit of the cBAF sub-family of mSWI/SNF chromatin remodeling enzymes, through unknown mechanisms. Unlike contributions to gene expression, it seems likely that this function is Pkm1-specific, and interrogation of the differences between Pkm1 and Pkm2 would be warranted to further address this function. A simple model would have Pkm1 directly phosphorylating Dpf2, but indirect mechanisms involving factors that control nuclear import or export are also possible.

In conclusion, the work here identifies novel, chromatin-based functions for Pkm2 during myoblast proliferation and differentiation and an unexpected role for Pkm1 in the nuclear localization of a chromatin remodeling enzyme needed for myoblast proliferation and differentiation. The findings increase the spectrum of Pkm functions that are distinct from the conventional role of Pkm enzymes in glycolysis and point to previously unappreciated roles for these enzymes in the regulation of the chromatin state and of gene expression.

## DATA AVAILABILITY STATEMENT

The datasets reported in this paper were deposited to GEO (Gene Expression Omnibus) under the accession number GSE252751.

## AUTHOR CONTRIBUTIONS

M. Olea-Flores, T. Padilla-Benavides, and A.N. Imbalzano conceptualized the work; T. Padilla-Benavides, P.R. Thompson, and A.N. Imbalzano acquired funding; M. Olea-Flores, T. Sharma, O. Verdejo-Torres, and I. DiBartolomeo performed experiments and acquired data; M. Olea-Flores, T. Sharma, P.R. Thompson, T. Padilla-Benavides, and A.N. Imbalzano analyzed data; M. Olea-Flores, T. Padilla-Benavides, and A.N. Imbalzano drafted and wrote the manuscript. All authors have reviewed and agreed to the submitted version of the manuscript.

## ACKNOWLEDGEMENTS

This work was supported by NIH grants R35 GM136392 to ANI, R01 AR077578 to TPB, and R35 GM118112 to PRT. We thank members of the Imbalzano and Padilla-Benavides lab for suggestions and comments on the manuscript.

