## Supplementary material for "Muscle-Specific Pyruvate Kinase Isoforms, Pkm1 and Pkm2, Regulate Mammalian SWI/SNF Proteins and Histone 3 Phosphorylation During Myoblast Differentiation": Supp. Figures & Supp. Tables 1 & 2

**FOR**

### Supplementary Figure 1

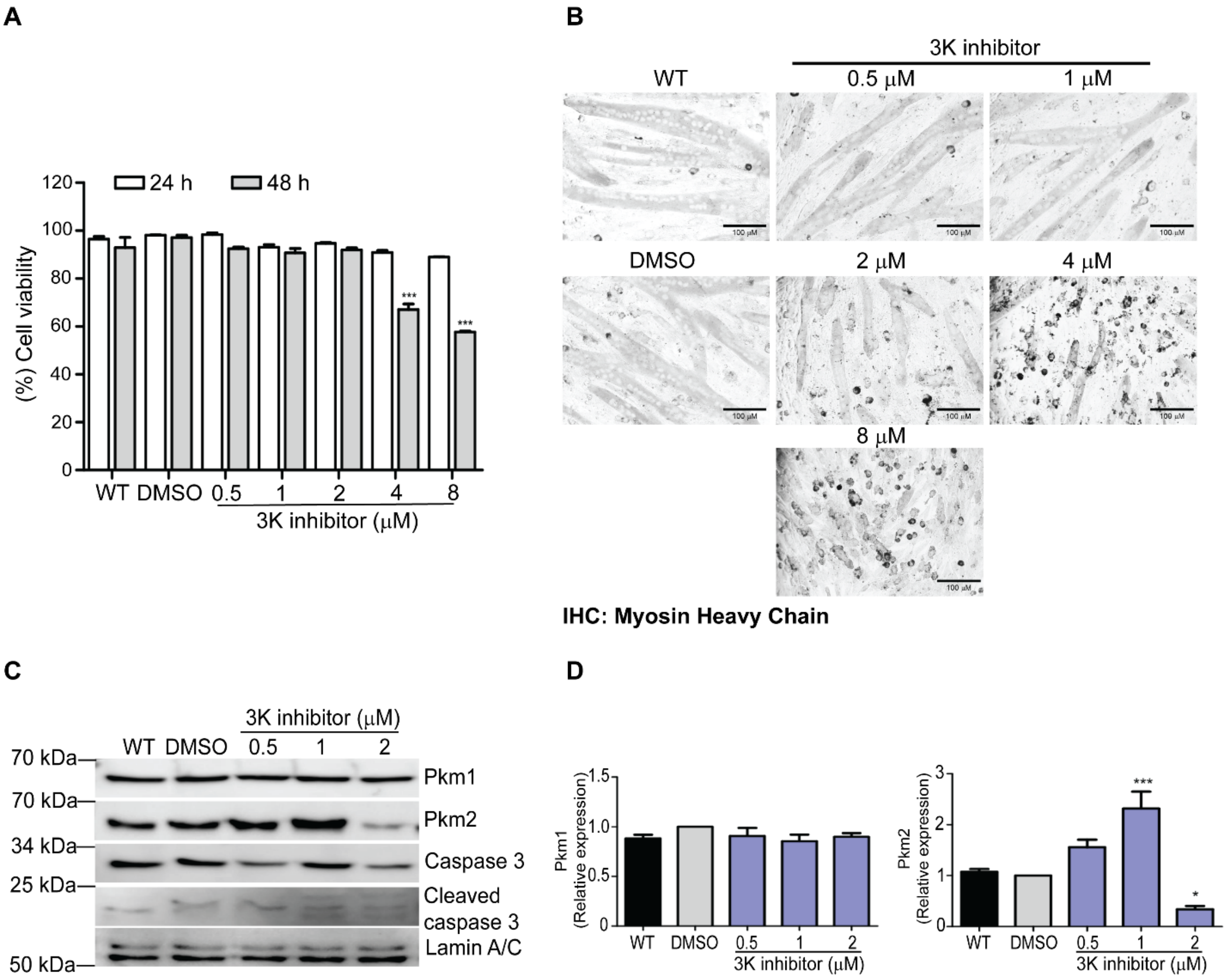


**Supplementary Figure 1. Titration of 3K. (A)** Cell viability was determined by the ratio of live to dead cells in the presence of concentrations from 0.5 to 8 μM of the 3K inhibitor. **(B)** Representative immunohistochemistry for Myosin Heavy Chain (MHC) in differentiating myoblasts in the presence of concentrations from 0.5 to 8 μM of the 3K inhibitor. **(C)** Representative western blots for the indicated proteins in the presence of the indicated concentrations of the 3K inhibitor. **(D)** Quantification of western blot results. Data in (A) and (D) are the mean ± SE of three independent biological replicates. *$P < 0.05$; **$P < 0.01$; ***$P < 0.001$. WT, wild type.

#### Supplementary Figure 2

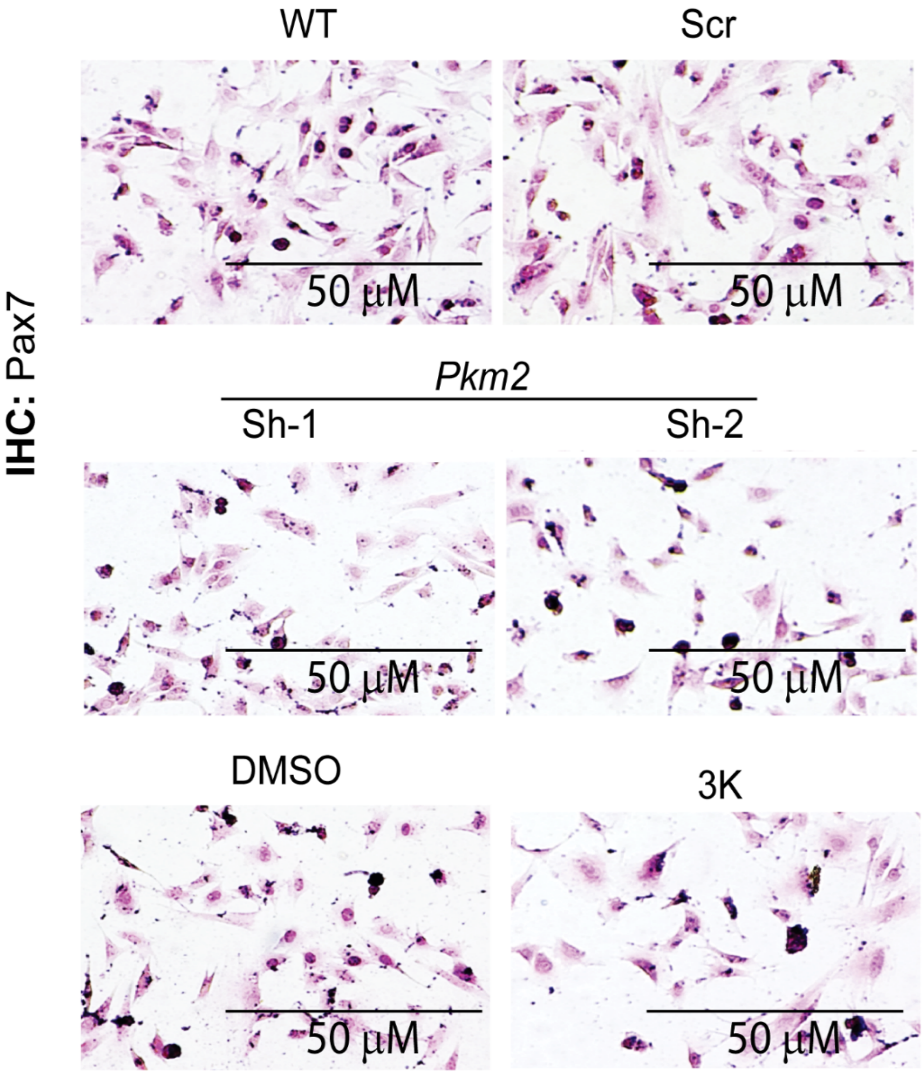


**Supplementary Figure 2. Cells subjected to *Pkm2* KD or Pkm2 inhibition showed reduced Pax7 staining.** Light micrographs of proliferating C2C12 myoblasts (controls, *Pkm2* KD, and in the presence of the 3K inhibitor) were cultured for 24 h and immunostained for Pax7. Representative images from three independent experiments are shown. WT, wild type, Scr, scrambled shRNA

### Supplementary Figure 3

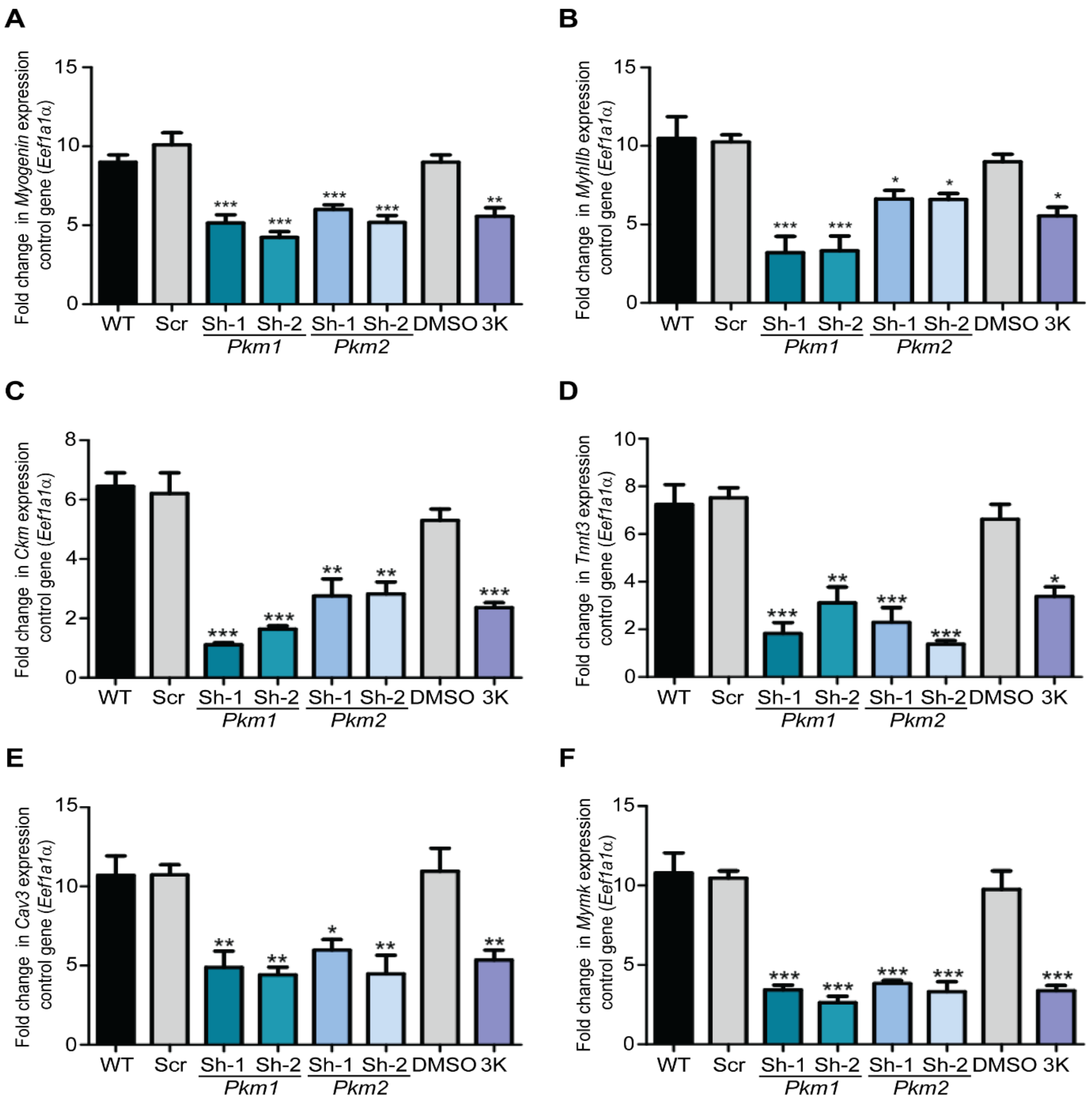


**Supplementary Figure 3. *Pkm2* KD or Pkm2 inhibition affected the expression of myogenic genes.** Steady state mRNA levels of (**A**) *Myogenin* (**B**) *MyhIIb*, (**C**) *Ckm,* (**D**) *Tnnt3,* (**E**) *Cav3, and* (**F**) *Mymk* determined by qRT-PCR from differentiating C2C12 myoblasts (WT, Scr, KD PKM1, KD PKM2 and in the presence of the 3K inhibitor). The data represent the mean ± SE from three independent biological experiments. *P < 0.05; **P < 0.01; ***P < 0.001. WT, wild type, Scr, scrambled shRNA.

### Supplementary Figure 4

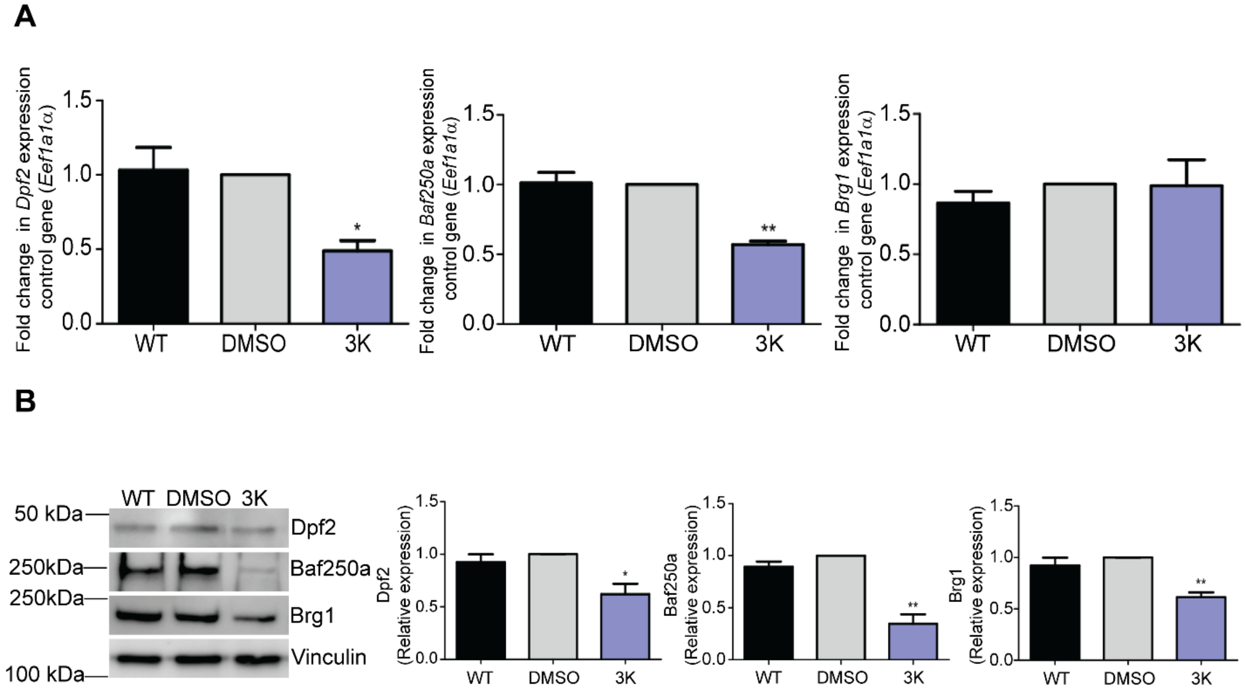


**Supplementary Figure 4. The Pkm2 inhibitor (3K), decreased the expression of Dpf2, Baf250a, and Brg1 in differentiating C2C12 myoblasts. (A)** mRNA levels of *Dpf2, Baf250a,* and *Brg1* in the presence or absence of Pkm2 inhibitor, 3K. **(B)** Representative immunoblots (left) and quantification (right) of Dpf2, Baf250a, and Brg1 protein levels in the presence and absence of the 3K Pkm2 inhibitor. Immunoblots against vinculin were used as loading controls. The data represent three independent biological experiments. Bar graphs show the mean ± SE. *P < 0.05, **P < 0.01 or ***P < 0.001. WT, wild-type.

### Supplementary Figure 5

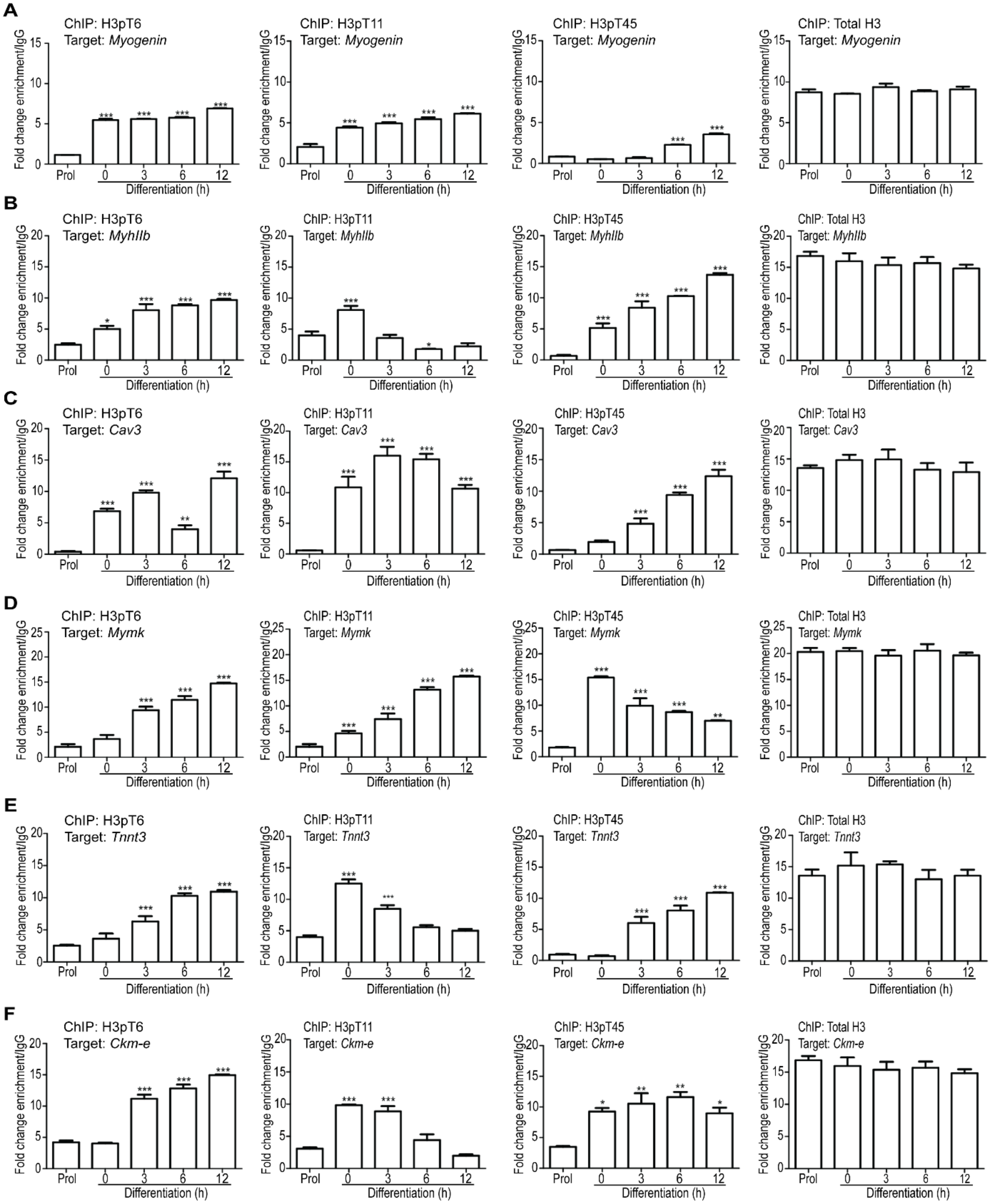

**Supplementary Figure 5. Incorporation of phosphorylated H3T6, H3T11 or H3T45 at myogenic promoters increased during differentiation.** ChIP-qPCR showing binding of phosphorylated H3T6, H3T11, H3T45 or total H3 to the **(A)** *Myogenin,* **(B)** *MyhIIb*, **(C)** *Cav3*, **(D)** Mymk **(E)***Tnnt3* promoters and **(F)** the *Ckm* enhancer (*Ckm-e)* in proliferating and differentiating C2C12 cells. The ChIP data was normalized using IgG as negative control of the ChIP. Bar graphs show the mean ± SE for three independent experiments. *P < 0.05; **P < 0.01; ***P < 0.001. Prol; proliferating myoblasts

### Supplementary Figure 6

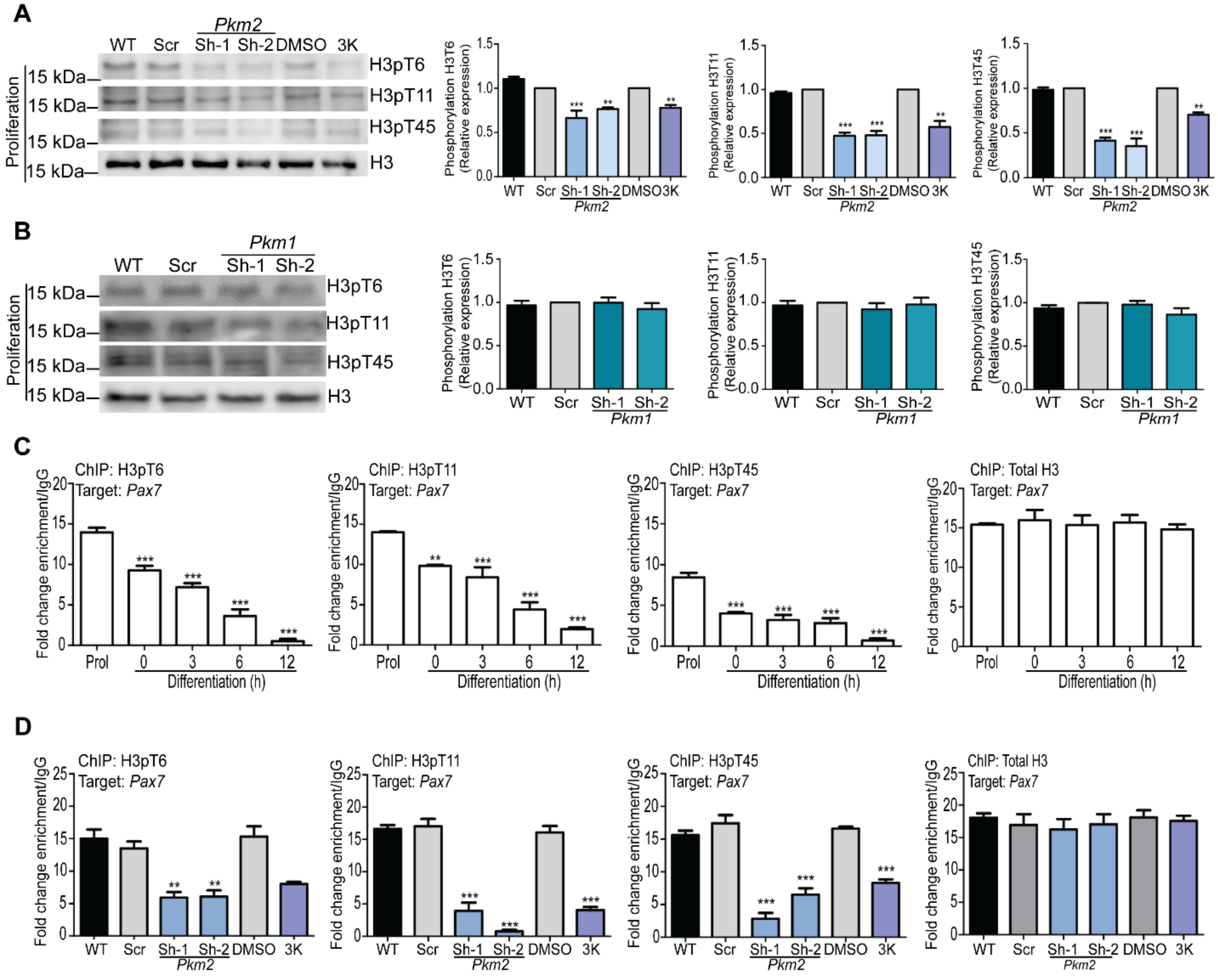


**Supplementary Figure 6. *Pkm2* KD decreased bulk intra-cellular levels and the incorporation into the *Pax7* promoter of phosphorylated H3T6, H3T11, and H3T45 in proliferating myoblasts*.*** **(A-B)** Representative western blots (left) and quantification (right) show phosphorylated H3T6, H3T11, and H3T45 levels in *Pkm2* KD myoblasts or in myoblasts in the presence of the 3K inhibitor **(A)** and in *Pkm1* KD myoblasts **(B)**. **(C)** ChIP-qPCR showing binding of phosphorylated H3T6, H3T11, H3T45, and total H3 to the *Pax7* promoter in proliferating myoblasts and myoblasts undergoing differentiation for up to 12 h. **(D)** ChIP-qPCR showing binding of phosphorylated H3T6, H3T11, H3T45, or total H3 to the *Pax7* promoter in *Pkm2* KD proliferating myoblasts or the presence of the 3K inhibitor. The ChIP data was normalized using IgG as a control for the ChIP. For all experiments, the data represent three independent biological experiments. Bar graphs show the mean ± SE. *P < 0.05; **P < 0.01; ***P < 0.001. WT, wild type, Scr, scrambled shRNA.
